# Quantitative Proteomics Links Mitochondrial Dysfunction to Metabolic Changes and Epithelial Differentiation Defects in Hyperoxia-Exposed Neonatal Airway Cells

**DOI:** 10.1101/2025.06.15.659571

**Authors:** Abhrajit Ganguly, Cynthia M. Carter, Aristides Rivera Negron, Hua Zhong, Alvaro Moreira, Matthew S. Walters, Lynette K. Rogers, Y.S. Prakash, Trent E. Tipple, Arlan Richardson

## Abstract

Premature infants often require supplemental oxygen therapy, however, exposure to supraphysiological oxygen (hyperoxia) can disrupt normal lung development and contribute to bronchopulmonary dysplasia (BPD). Mitochondrial dysfunction is increasingly recognized as a contributor to hyperoxia-induced BPD. However, the effects of hyperoxia on mitochondrial function and mucociliary differentiation in the developing upper airway epithelium remain poorly understood. This study tested the hypothesis that hyperoxia impairs neonatal airway mucociliary differentiation by disrupting mitochondrial bioenergetic function. Neonatal tracheal airway epithelial cells (nTAECs) from term infants (n=5) were cultured in a 3D air-liquid interface (ALI) model and exposed to 60% O₂ during the mid-phase of differentiation (ALI day 7-14). Cellular phenotype was assessed using immunofluorescence staining and gene expression analyses. Mitochondrial function was evaluated through Seahorse metabolic flux analysis, and global protein changes were characterized by quantitative proteomics. Hyperoxia exposure significantly impaired terminal epithelial differentiation, characterized by reductions in ciliated and goblet cells. Seahorse assay revealed a decrease in baseline oxygen consumption and mitochondrial ATP production, accompanied by a compensatory increase in glycolytic ATP production. Quantitative proteomics identified disruption of mitochondrial Complex I as a central feature of the hyperoxic response. Downstream proteomic pathway analyses further confirmed the metabolic shift from mitochondrial to glycolytic ATP production and demonstrated altered epithelial differentiation pathways, including NOTCH and TGF-β signaling. These findings reveal that moderate hyperoxia impairs mitochondrial bioenergetics and alters metabolic programming, leading to disrupted mucociliary differentiation. Future *in vivo* studies should evaluate mitochondrial oxidative fitness as a therapeutic target in neonatal lung disease.

**NEW & NOTEWORTHY:** We report that moderate hyperoxia during a critical window of mucociliary differentiation disrupts terminal maturation in neonatal airway epithelial cells cultured in a 3D model. Hyperoxia induced mitochondrial bioenergetic dysfunction and metabolic reprogramming, with proteomic analysis identifying Complex I disruption as a key driver of impaired differentiation. Overall, these findings reveal a previously underrecognized link between mitochondrial bioenergetics and airway epithelial development, positioning metabolic dysfunction as an early trigger of hyperoxia-induced neonatal airway injury.

## INTRODUCTION

Globally, an estimated 13.5 million babies are born preterm (<37 weeks or wks) every year, of which over 2 million are born very preterm (<32 wks)(1). Of the very preterm infants who survive the first years of life, many suffer from significant lung morbidity with increased wheezing disorders, exercise limitation and recurrent infections leading to significant healthcare utilization and cost (2–8). The disease burden further rises depending on the degree of prematurity and whether the infant develops bronchopulmonary dysplasia (BPD), a form of heterogenous chronic lung disease uniquely seen in premature infants (9–14). Supraphysiological oxygen (O_2_) therapy or hyperoxia is commonly used in the care of premature infants (9, 10, 15) and contributes to the development of BPD. Over the past decade, most of the research on hyperoxia induced BPD has focused on alveolar, airway smooth muscle and lung endothelial development (16–19). Despite the mucociliary epithelium being in the direct contact with oxidative stress from elevated oxygen exposure, the impact of hyperoxia on upper airway epithelial development and function remains understudied.

Proper functioning of the airway epithelium is essential for effective mucociliary clearance, defense against pathogens and allergens, and overall airway development(20). Human lung development progresses through several distinct phases, beginning with the embryonic phase *in utero* to the alveolar phase, which extends into adolescence (21). The early airway epithelium is initially comprised of undifferentiated columnar cells which give rise to early airway epithelial progenitors (20–23). Epithelial differentiation in humans begins around 14 to 16 wks of gestation in the early canalicular phase (16 to 26 wks gestation) with the emergence of club cells, followed by the appearance of ciliated and mucus forming goblet cells, each with specialized characteristics and function (20). Epithelial differentiation continues through the saccular phase (26 to 36 wks gestation) and by the early alveolar phase (> 36 weeks), the mucociliary epithelium is structurally and functionally mature. In the current context of premature birth, most preterm babies are delivered during late canalicular to saccular phase (∼22 to 36 wks) of lung development, a critical window during which epithelial maturation is actively underway (10). Oxygen therapy and hyperoxia exposure in preterm infants typically occur during this sensitive period, when airway epithelial differentiation and function are still developing (21). In extremely preterm infants (born before 28 wks gestation), the lungs contain very few mature alveoli at birth, making the immature airway epithelium the primary site of oxidative injury.

As the hub of cellular redox metabolism, mitochondrial dysfunction has been implicated in the pathogenesis of oxidative lung diseases like BPD and chronic obstructive pulmonary disease (COPD) (24). Cellular differentiation, including in the airway mucociliary epithelium, relies on key metabolic switches regulated by efficient mitochondrial bioenergetic function (25–27). A recent study showed that even moderate prematurity (28-29 wks gestation) causes airway epithelial mitochondrial dysfunction, and abnormal ciliary development (28). Animal studies with hyperoxia-induced BPD have demonstrated that hyperoxia significantly inhibits mitochondrial bioenergetic machinery, especially Complex I (29), and attenuates oxidative phosphorylation resulting in hypo-alveolarization (30). In the same study, use of a Complex I inhibitor reproduced the aberrant lung phenotype, highlighting the critical role of intact Complex I and bioenergetic function during lung development. However, the contribution of mitochondrial bioenergetics to neonatal airway mucociliary development and function remain poorly understood. In a recent study, we utilized a human neonatal patient sample-derived tracheal airway epithelial cells (nTAECs) in a novel translational 3D culture model to show that 60% O_2_ exposure during mid-phase of mucociliary differentiation induces significant cellular oxidative stress (31). Building on these findings, we hypothesized that 60% O_2_ exposure in nTAECs during mid-phase of differentiation disrupts normal airway epithelial maturation by impairing mitochondrial bioenergetic function. The overall objectives of this study were to, a) characterize the neonatal airway epithelial phenotype following hyperoxia exposure; b) assess the functional consequences of hyperoxia exposure on bioenergetic profile of nTAECs; and c) utilize quantitative proteomics to define hyperoxia-sensitive molecular networks in the developing nTAECs, focusing on mitochondrial and oxidative stress pathways.

## MATERIALS AND METHODS

### Neonatal Tracheal Aspirate Sample Collection and Processing

Tracheal aspirate samples were collected from neonates admitted in the NICU after informed consent from parents, and the protocol used for collection, transport, and storage has been approved by the Institutional Review Board (IRB) of the University of Oklahoma Health Sciences Center (IRB 14377). For this study, patient samples were collected from term (≥ 37 wks gestation) intubated infants admitted in the NICU, during routine suction of the endotracheal tube. All samples were de-identified before transport to the lab except for gestational age at birth, sex and primary diagnosis at admission to the NICU. A table with the clinical characteristics of the infants are included in the supplementary material (Supplemental table 1). This study was not powered to detect differences by sex owing to small sample size. Fresh tracheal aspirate samples were collected and processed as described previously (31). In short, approximately 1ml of tracheal aspirate sample was collected in a mucus trap under sterile condition and kept on ice for transport to the lab. Following transport to the lab, the sample was diluted with sterile PBS, centrifuged at 250 rcf for 5 mins at 20 - 23 °C and the supernatant (containing mucus) was discarded.

### Isolation and Expansion of Neonatal Tracheal Airway Epithelial Cells (nTAECs)

After initial processing and removal of mucus, the samples were resuspended in 5ml of BronchiaLife^TM^ Epithelial Airway Medium or BLEAM (LifeLine cell technology, MD, USA), plated on a 804G conditioned media coated T25 flask (Passage 0 or P0) and placed in the incubator at 37 °C, 5% CO_2_ (31). Media was changed 24 hrs after plating to wash out unattached cells and subsequently every 48 hrs. Cuboidal-shaped cells appear 7-10 days post-plating. By around 3 weeks post-plating, the cells are densely packed and require trypsinization for subsequent passaging, expansion, and storage. Early passage (P1 - P2) cells were placed in cryotubes (2.5 x 10^6^ cells per 500 µL of freezing media per cryotube) and stored in liquid nitrogen for long-term storage. nTAECs were then utilized in 3D culture as described below for differentiation and submerged culture for seahorse experiments.

### Air-liquid Interface (ALI) Culture and Hyperoxia Exposure

Frozen nTAECs were rapidly thawed from liquid nitrogen, cells were counted to determine the total and live cell number, plated at 2.1 x 10^6^ cells in 15ml of cell suspension in a T75 flask and incubated at 37 °C, 5% CO_2_. Media was changed 24 hrs after plating to wash out unattached cells and subsequently every 48 hrs. At 80-90% confluence, cells were trypsinized and counted to determine the total and live cell number. 6.5mm Transwell^®^ with 0.4 μm pore polyester membrane inserts (Corning, NY, USA) were coated with 804G conditioned media. 1 x 10^5^ cells in 100 μL cell suspension was added to each apical (upper) chamber of the ALI inserts and 1ml BLEAM media was added to the basolateral (lower) chamber and incubated at 37 °C, 5% CO_2_ (ALI day −2). Media was replaced in both apical and basolateral chamber every 24hrs. Once the cells reached confluence on the ALI insert membrane (after 48 hrs), media in the basolateral chamber was exchanged with ALI epithelial differentiation media (LifeLine cell technology, MD, USA) and the apical chamber was left media-free and exposed to air (ALI day 0). ALI differentiation was continued for 14 days (ALI day 14). Media in the basolateral chamber was exchanged every 48 hrs with new differentiation media and any leakage into the apical chamber was aspirated every 24hrs for the initial 5-7 days of ALI culture. On ALI day 7, ALI inserts assigned in the hyperoxia group were incubated in the TriGas incubator (Thermo Fisher Scientific, MA, USA) under 60% O_2_ exposure and continued for 7 days until ALI day 14. Control inserts were incubated at 37 °C, 5% CO_2_ (room air or normoxia). Media in the basolateral chamber was exchanged every 48 hrs with new differentiation media.

### Trans-Epithelial Electrical Resistance (TEER) measurement

TEER was measured using Epithelial Volt-Ohm Meter or EVOM (World Precision Instruments, FL, USA) to assess epithelial barrier function and cell-layer integrity as previously described (31). Background resistance value (in Ohm or Ω) was measured using an 804G conditioned media-coated ALI insert (devoid of any cells), subtracted from the test insert resistance value and the difference was multiplied by the cell growth surface area to obtain TEER value for each test insert. TEER values were calculated in hyperoxia exposed and control nTAECs on ALI day 14 and values were expressed normalized to room air control for each corresponding donor cells.

### Immunofluorescent Staining and Quantification of Epithelial Differentiation

Cells cultured at air-liquid interface (ALI) were fixed in situ using 10% neutral buffered formalin (NBF) for 20 minutes at room temperature. After fixation, the wells were rinsed with phosphate-buffered saline (PBS) and stored at 4°C until staining. For immunostaining, cells were permeabilized with 0.1% Triton X-100 (Cat# 194854, MP Biomedicals, Irvine, CA), followed by blocking in 5% donkey serum to reduce nonspecific binding. Samples were then incubated for 2 hours at room temperature with primary antibodies targeting key epithelial cell markers: P63α (basal cells; 1:100, Cat# 13109S, Cell Signaling Technology), Secretoglobin 1A1 or SCGB1A1 (club cells; 1:200, Cat# RD181022220-01, BioVendor LLC), Mucin-5AC or MUC5AC (goblet cells; 1:100, Cat# MA5-12178, Thermo Scientific), and acetylated tubulin or ACTUB (ciliated cells; 1:200, Cat# T7451, Sigma-Aldrich). Following primary antibody incubation, cells were washed thoroughly with PBS and then incubated with appropriate fluorescently conjugated secondary antibodies for 45 minutes at room temperature (1:200 dilution; Donkey anti-mouse Alexa Fluor 594 and Donkey anti-rabbit Alexa Fluor 647, Cat# 715-586-151, Jackson ImmunoResearch). Nuclear counterstaining was performed using DAPI. The Transwell® membranes were carefully excised and mounted onto glass slides using ProLong™ Gold Antifade Mountant without DAPI (Cat# P36930, Thermo Scientific). Fluorescent images were acquired using an Olympus BX43 upright microscope (Olympus Corporation). For each sample, three to four randomly selected fields were imaged, and a minimum of 1500 epithelial cells (DAPI stained nuclei) were analyzed. Positive staining for each marker was quantified using ImageJ software (version 2.160/1.54g, NIH), and results were normalized to the total number of DAPI-stained nuclei per image.

### RNA Extraction and Gene Expression Analysis

Total RNA was isolated directly from cells in ALI wells using the PureLink™ RNA Mini Kit (Cat# 12183018A, Thermo Scientific, Waltham, MA), following the manufacturer’s protocol. To eliminate residual genomic DNA, on-column DNase digestion was performed using the PureLink™ DNase Set (Cat# 12185-010, Thermo Scientific). Complementary DNA (cDNA) was synthesized from equal amounts of RNA (50–250 ng per sample) using random hexamer primers provided in the High-Capacity cDNA Reverse Transcription Kit (Cat# 4374966, Thermo Scientific, Applied Biosystems™). Quantitative real-time PCR (qRT-PCR) was carried out using TaqMan™ Fast Advanced Master Mix (Thermo Scientific) on a QuantStudio 3 Real-Time PCR system (Applied Biosystems). Gene expression was analyzed using the comparative cycle threshold (ΔCt) method, with GAPDH (Hs02786624_g1) serving as the endogenous control. TaqMan™ gene expression assays were used for target detection - KRT5 (Hs00361185_m1), FOXJ1 (Hs00230964_m1), SCGB1A1 (Hs00171092_m1), MUC5AC (Hs01365616_m1) following the manufacturer’s recommended thermal cycling parameters. Gene expression was measured in three technical replicates per condition for each donor cells. The average of these three technical replicates was used to represent the gene expression value for that sample.

### Cellular bioenergetic profiling with Seahorse

To assess mitochondrial and glycolytic ATP production rates, the Seahorse XFp Real-Time ATP Rate Assay Kit (Agilent, Cat# 103591-100) was used according to the manufacturer’s instructions with the Agilent Seahorse XFp Analyzer (Agilent Technologies, Santa Clara, CA, USA). nTAECs (n = 5 donor cells) were cultured under submerged conditions and exposed to either room air (21% O₂) or 60% O₂ for 48 hours. On the day of the assay, real-time oxygen consumption rate (OCR) and extracellular acidification rate (ECAR) were recorded under basal conditions and after sequential injection of oligomycin (final concentration 1.5 µM), and a mixture of rotenone and antimycin A (final concentration 0.5 µM each). The resulting data were analyzed using the Agilent XF Real-Time ATP Rate Assay Report Generator to quantify mitochondrial ATP production (mitoATP), glycolytic ATP production (glycoATP), and total cellular ATP production.

### Data Independent Acquisition (DIA) Proteomics

DIA Proteomics was performed on ALI day 14 following hyperoxia exposure for 7 days (ALI day 7 to 14) utilizing whole cell lysates. Total protein from each sample (n = 5 donor cells per group) was reduced, alkylated, and purified by chloroform/methanol extraction prior to digestion with sequencing grade modified porcine trypsin (Promega). Tryptic peptides were then separated by reverse phase XSelect CSH C18 2.5 um resin (Waters) on an in-line 150 x 0.075 mm column using an UltiMate 3000 RSLCnano system (Thermo Scientific). Peptides were eluted using a 60 min gradient from 98:2 to 65:35 buffer A:B ratio (Buffer A = 0.1% formic acid, 0.5% acetonitrile, Buffer B = 0.1% formic acid, 99.9% acetonitrile). Eluted peptides were ionized by electrospray (2.2 kV) followed by mass spectrometric (MS) analysis on an Orbitrap Exploris 480 mass spectrometer (Thermo Scientific). To assemble a chromatogram library, six gas-phase fractions were acquired on the Orbitrap Exploris with 4 m/z DIA spectra (4 m/z precursor isolation windows at 30,000 resolution, normalized AGC target 100%, maximum inject time 66 ms) using a staggered window pattern from narrow mass ranges using optimized window placements. Precursor spectra were acquired after each DIA duty cycle, spanning the m/z range of the gas-phase fraction (i.e. 496-602 m/z, 60,000 resolution, normalized AGC target 100%, maximum injection time 50 ms). For wide-window acquisitions, the Orbitrap Exploris was configured to acquire a precursor scan (385-1015 m/z, 60,000 resolution, normalized AGC target 100%, maximum injection time 50 ms) followed by 50x 12 m/z DIA spectra (12 m/z precursor isolation windows at 15,000 resolution, normalized AGC target 100%, maximum injection time 33 ms) using a staggered window pattern with optimized window placements. Precursor spectra were acquired after each DIA duty cycle. Following MS acquisition, data were searched using an empirically corrected library against the UniProt *Homo sapien* database and a quantitative analysis was performed to obtain a comprehensive proteomic profile. Proteins were identified and quantified using EncyclopeDIA (32) and visualized with Scaffold DIA using 1% false discovery thresholds at both the protein and peptide level. Protein MS2 exclusive intensity values were assessed for quality using ProteiNorm (33). The data was normalized using VSN (34) and analyzed using proteoDA to perform statistical analysis using Linear Models for Microarray Data (limma) with empirical Bayes (eBayes) smoothing to the standard errors (35, 36). Proteins with an adjusted p-value (FDR) < 0.05 and absolute fold change > 2 (log2FC > 1) were defined as significantly differentially expressed. The mass spectrometry proteomics data have been deposited to the ProteomeXchange Consortium via the PRIDE (37) partner repository with the dataset identifier PXD064170 and 10.6019/PXD064170.

### Downstream Visualization and Analysis of Proteomic Data

Principal Component Analysis (PCA) was performed to visualize global proteomic differences between room air and hyperoxia-exposed nTAECs. Raw protein abundance values from DIA proteomics were first log2-transformed with a pseudocount of 1 to reduce skewness and stabilize variance. Proteins with greater than 50% missing values across all samples were excluded. Remaining missing values were imputed using the mean of each protein across all samples. For genes with duplicate entries, mean expression was calculated to consolidate them into a single identifier. Zero-variance proteins were removed prior to PCA to avoid artifacts during scaling. PCA was performed using the prcomp() function in R with centering and unit variance scaling enabled. The first two principal components were plotted, and samples were grouped by exposure condition. To visualize the distribution of biological replicates without assuming parametric shape, convex hulls were drawn around sample groups utilizing the ggforce package. This approach highlights the spatial clustering of samples within each experimental condition and emphasizes differences in overall proteomic profiles between room air and hyperoxia groups. Volcano plot was generated in R (version 4.4.0) using ggplot2(38) and ggrepel(39). A relaxed threshold of |log2FC| > 0.25 and FDR < 0.05 was applied for plotting. Heatmaps were generated using the pheatmap package(40). The top differentially expressed proteins (DEPs) were selected, log2-transformed, and z-score normalized across rows. Rows (proteins) were clustered hierarchically using Euclidean distance and complete linkage. To gain functional insights into the differentially expressed proteins, pathway enrichment analyses were performed using both publicly available and proprietary databases(41). Gene ontology enrichment for Biological Processes (GO BP), as well as pathway enrichment using Reactome was conducted in R using the clusterProfiler(42) and ReactomePA(43) packages. Protein identifiers were mapped to gene symbols, and enrichment analyses were run using a background of all detected proteins. Terms with a Benjamini-Hochberg adjusted p-value < 0.05 were considered significantly enriched. Additionally, canonical pathway and biological process analyses were performed using Ingenuity Pathway Analysis (IPA, QIAGEN Inc.)(44) based on differentially expressed protein identifiers and their associated expression fold changes. IPA identified enriched canonical pathways and biological functions using right-tailed Fisher’s exact test and predicted pathway activation states using z-scores. Significant IPA terms were filtered based on -log(p-value) > 1.3 (corresponding to p < 0.05) and absolute z-score > 2. Gene Set Enrichment Analysis (GSEA) was performed to identify coordinated changes in predefined biological pathways across the full ranked proteome. Proteins were ranked by their signed log10-transformed p-values multiplied by the direction of fold change. GSEA was carried out using the clusterProfiler package in R with gene sets derived from the Molecular Signatures Database (MSigDB) WikiPathway and Hallmark pathway(45), as well as the Reactome database via the ReactomePA package. Enrichment scores (ES) and normalized enrichment scores (NES) were calculated based on 1,000 permutations. Gene sets with a false discovery rate (FDR) q-value < 0.25 and nominal p-value < 0.05 were considered significantly enriched. Enrichment results from these approaches were visualized using dot plots and heatmaps generated in R with the ggplot2, enrichplot, and ComplexHeatmap packages.

### Statistical methods

Statistical analysis was performed using GraphPad Prism 10.4.1 (GraphPad Software, CA, USA) software. For most analyses, fold changes were determined relative to expression or changes in room air controls. Given the small sample size, non-parametric statistical tests were applied for statistical analysis as appropriate and are indicated in figure legends. A *p*-value of ≤0.05 was considered statistically significant. Analysis of proteomic data are described under the DIA Proteomics method section. A comprehensive list of key proteins referenced throughout the study, including those identified by DIA proteomics is provided in Supplemental Table 2, along with their gene symbols, full protein names, and associated biological pathways.

## RESULTS

### Patient sample-derived nTAECs in ALI culture undergo mucociliary differentiation into mature epithelial subtypes over 14 days

We postulated that nTAECs undergo terminal differentiation by ALI day 14. To test this, basal nTAECs were isolated and cultured from neonatal tracheal aspirate samples (Fig 1A) and subsequently cultured in ALI over 14 days to facilitate mucociliary differentiation (Fig 1B). To evaluate mucociliary differentiation, we performed gene expression analysis and immunofluorescence staining of nTAECs utilizing lineage-specific epithelial markers for basal (KRT5 and P63-α), club (SCGB1A1), goblet (MUC5AC) and ciliated (FOXJ1 and ACTUB) cells. By ALI day 14, gene expression analysis revealed a significant increase in SCGB1A1 (p=0.028), MUC5AC (p=0.028) and FOXJ1 (p=0.028) expression compared to ALI day 0 (Fig. 1C), suggesting progression toward differentiated cell types. On ALI day 0, IF staining showed nearly all cells were positive for the basal cell marker P63-α, indicating an undifferentiated basal cell phenotype (Fig 1D). IF staining at ALI day 14 further validated these findings, showing the presence of SCGB1A1+ club cells, ACTUB+ ciliated cells, and MUC5AC+ goblet cells (Fig. 1E), confirming terminal mucociliary differentiation of nTAECs.

**Figure 1.**
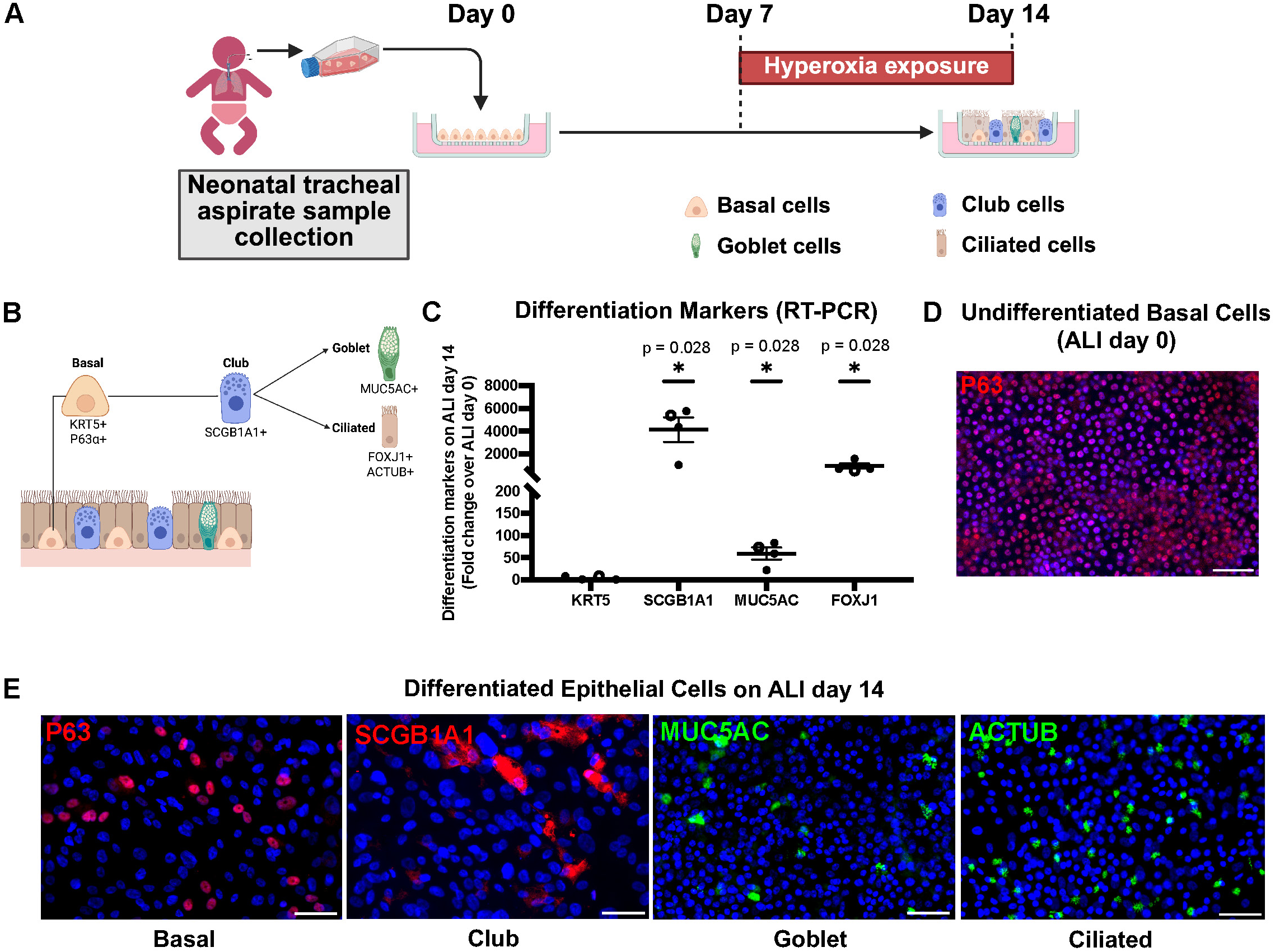
Neonatal tracheal aspirate-derived airway epithelial cells (nTAECs) undergo terminal mucociliary differentiation during 3D air-liquid interface (ALI) culture. (A) Experimental timeline for neonatal tracheal aspirate collection to ALI culture of nTAECs with hyperoxia exposure. (B) Normal airway mucociliary differentiation from basal cells. (C) Quantitative PCR analysis of epithelial cell-specific markers for basal (KRT5), club (SCGB1A1), ciliated (FOXJ1), and goblet (MUC5AC) cells on ALI day 14 compared to ALI day 0. Relative gene expression was normalized to GAPDH (endogenous control) and plotted as fold change over ALI day 0. Data presented as mean (n = 4 donor samples. Solid black circles represent males; open circle represents female). Error bars indicate SEM. Statistical analysis was performed using Mann-Whitney U test. (D-E) Immunofluorescent staining with representative images of airway progenitor (P63⍺, red) performed on ALI day 0 and 14, and club (SCGB1A1, red), goblet (MUC5AC, green) ciliated (ACTUB, green) cells were performed on ALI day 14. Nuclei are counterstained blue with DAPI. Scale bar = 50 µm.

### Hyperoxia exposure disrupts normal epithelial differentiation of nTAECs in ALI culture

Hyperoxia exposure with 60% O_2_ was initiated at ALI day 7 after the formation of a mature cell monolayer as we had shown in previously published work from our group(31). We hypothesized that moderate hyperoxia (60% O_2_) exposure during mid-phase of mucociliary differentiation (ALI day 7 to 14) would disrupt normal epithelial maturation. To test our hypothesis, we performed IF staining with lineage specific epithelial markers and compared the room air and hyperoxia groups. Hyperoxia exposure significantly decreased the number of ACTUB+ ciliated cells (p=0.008) and MUC5AC+ goblet cells (p=0.007) compared to room air control (Fig 2A). There was a slight increase in P63α+ basal cells (p=0.12) and SCGB1A1+ club cells (p=0.682) in the hyperoxia-exposed group. Together, these findings suggest that hyperoxia impairs terminal differentiation of nTAECs, favoring a progenitor-like state characterized by persistence of basal and club cells and reduced emergence of mature ciliated and goblet cells. To determine whether these differentiation changes were accompanied by alterations in epithelial barrier integrity, we measured TEER values on ALI day 14. TEER measurements did not differ significantly between hyperoxia and room air groups (p>0.999) (Fig. 2B), indicating that barrier function remained intact despite impaired differentiation.

**Figure 2.**
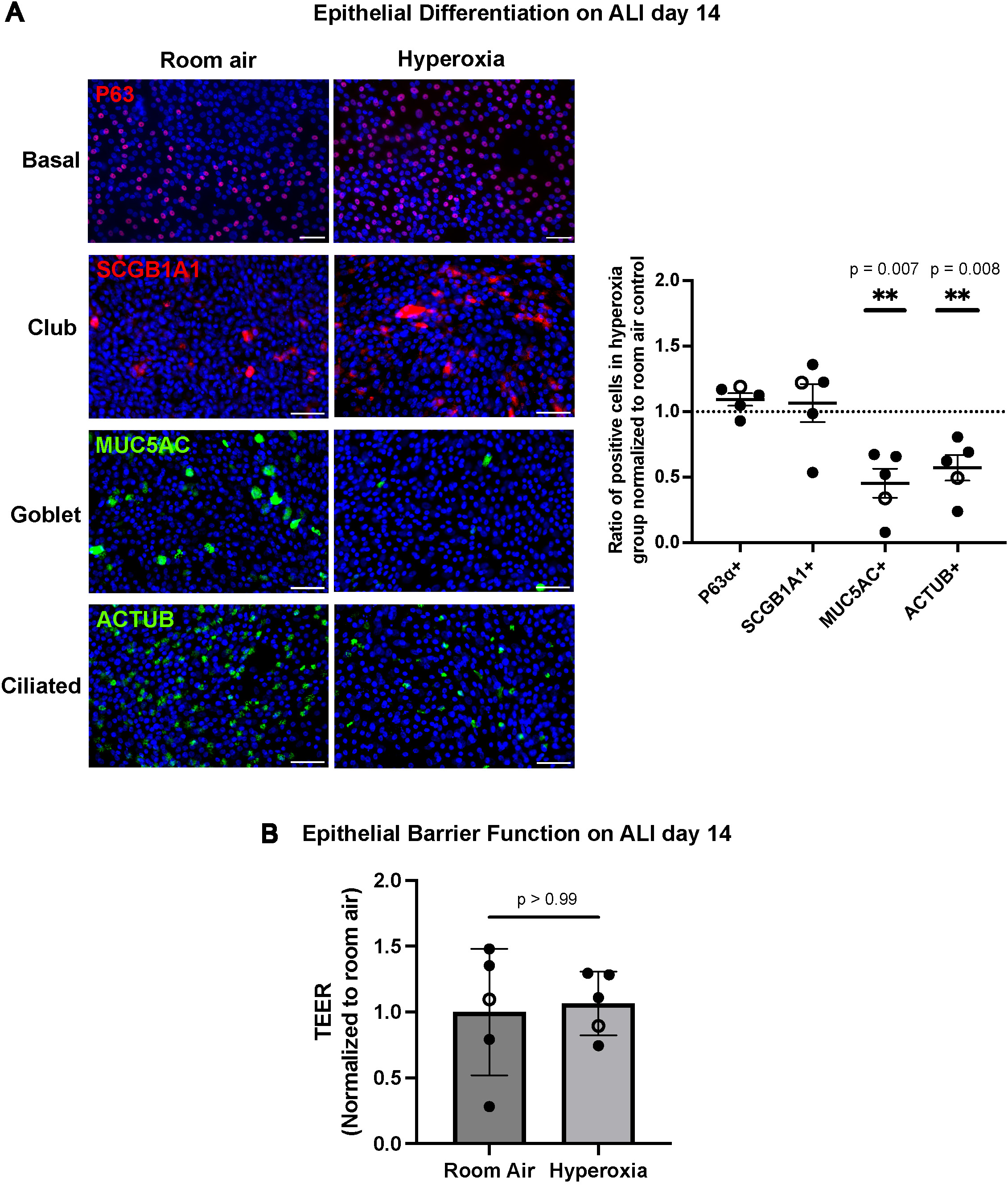
Hyperoxia exposure disrupts normal epithelial differentiation in nTAECs with decreased number of ciliated and goblet cells. (A) Immunofluorescent staining of airway progenitor (P63⍺, red), ciliated (ACTUB, green), club (SCGB1A1, red) and goblet (MUC5AC, green) cells were performed on ALI 14 following room air or hyperoxia exposure. Nuclei are counterstained blue with DAPI. Each data points represent the ratio of positive cells for each donor (n = 6 donor sample. Solid black circles represent males; open circle represent female) normalized to room air control. Statistical analysis was performed Mann-Whitney U test. Scale bar = 50 µm. (B) TEER measurement on ALI day 14 comparing room air and hyperoxia exposed nTAECs do not show any significant difference. Data points represent average TEER values for each donor (n=5 donors, 3-4 wells per donor. Solid black circles represent males; open circle represent female). Statistical analysis was performed Mann-Whitney U test.

### Mitochondrial metabolic flux analysis shows hyperoxia exposure results in alteration of cellular bioenergetic profile in nTAECs

Cellular differentiation, including the developing airway epithelium, is an energy-intensive process and thus requires a robust mitochondrial network and intact cellular bioenergetic machinery (25, 27, 46). Neonatal hyperoxia exposure has been shown to cause mitochondrial dysfunction leading to alveolarization defects in an animal model of BPD (30, 47). We hypothesized that moderate hyperoxia exposure will alter the bioenergetic profile of nTAECs. To test this hypothesis, we performed Seahorse metabolic flux analysis to investigate the effect of hyperoxia on cellular bioenergetic profile of nTAECs. Due to technical limitations in performing the assay on transwell inserts with mature monolayers (ALI day 14), we utilized undifferentiated basal nTAECs in submerged culture. OCR and ECAR were quantified following sequential addition of oligomycin and rotenone/antimycin A (Rot/AA). OCR profiles differed markedly between hyperoxia-exposed and normoxia control cells (Fig 3A). Hyperoxia-exposed cells showed reduced OCR at baseline which further dropped following oligomycin injection in some donor cells, suggesting lower ATP-linked or mitochondrial respiration following hyperoxia. Non-mitochondrial respiration, which was calculated by OCR post-Rot/AA injection, did not show marked alteration in the hyperoxia group. Impact of hyperoxia on ECAR responses to hyperoxia varied among donors, with some showing marked increases, while others showed minimal change (Fig 3B). Quantification of ATP sources revealed that hyperoxia significantly reduced (p=0.01) mitochondrial-derived ATP (mitoATP) and increased (p=0.01) glycolytic ATP (glycoATP) compared to room air controls (Fig 3C).

**Figure 3.**
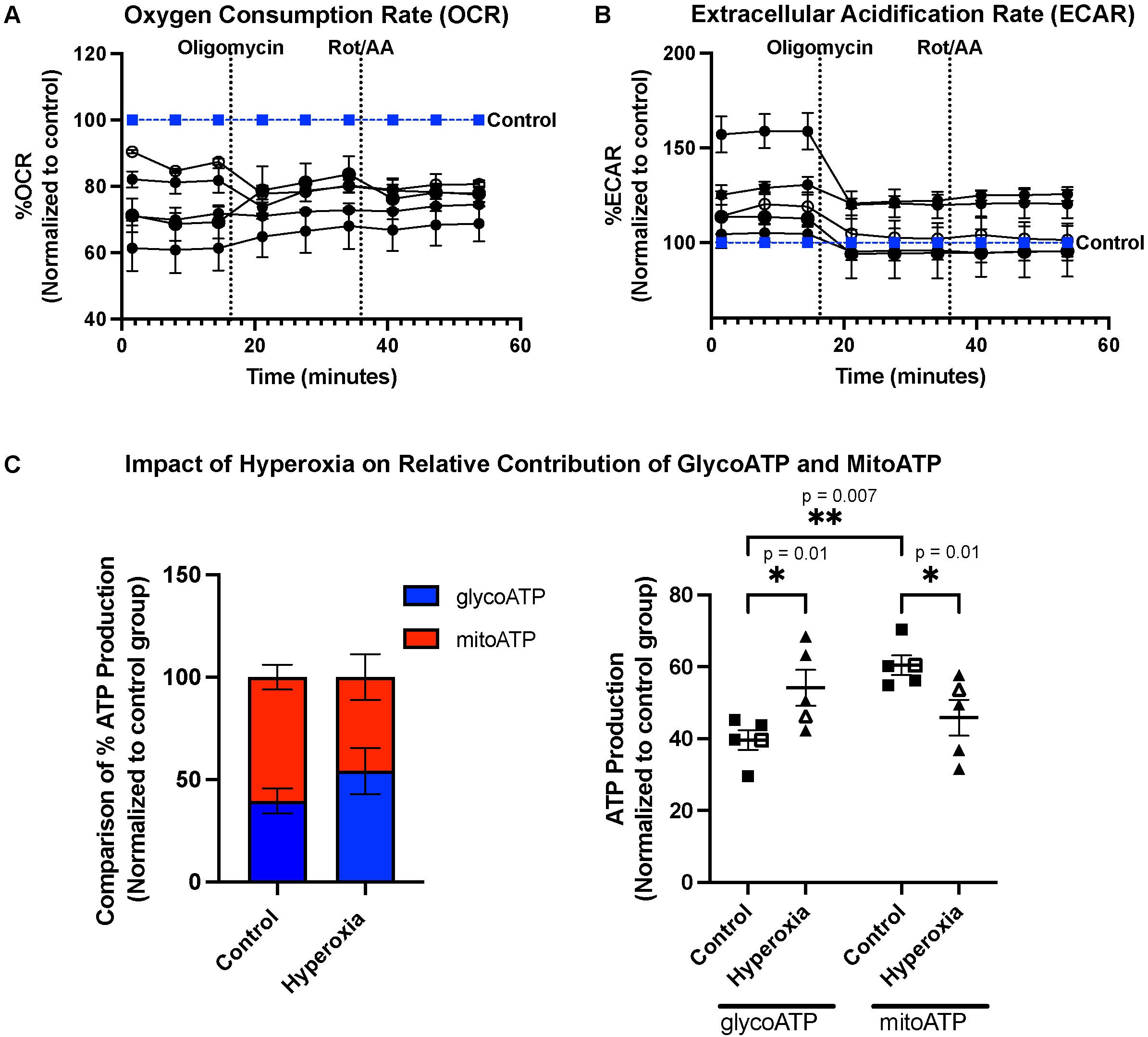
Hyperoxia impairs the cellular bioenergetic profile of neonatal airway epithelial cells and alters the metabolic phenotype with significantly reduced mitochondrial ATP production and increased glucose metabolism. (A) Oxygen consumption rate (OCR) and Extracellular Acidification Rate (ECAR) was measured in nTAECs using a Seahorse XF Analyzer following hyperoxia exposure. Cells were sequentially treated with oligomycin and rotenone/antimycin A (Rot/AA) to assess ATP production. Data are expressed as percent OCR or ECAR normalized to room air controls with mean ± SD (n = 5 donor cells. Blue line and squares represent normalized combined room air controls; Solid black circles represent hyperoxia exposed male donor cells; open circle represent hyperoxia-exposed female donor cells) (B) Total ATP production rate was calculated, normalized and partitioned into mitochondrial (mitoATP, red) and glycolytic (glycoATP, blue) contributions using the Seahorse ATP Rate Assay, n=5 donor samples. (C) Impact of hyperoxia on mitochondrial (mitoATP) vs glycolytic (GlycoATP) ATP production compared to control. Data are presented as mean ± SEM (n=5 donor samples. Solid black squares and triangles represent males; open squares and triangles represent female). Statistical analysis was performed using Two-way ANOVA with Šídák’s multiple comparison test.

### Quantitative proteomics reveals hyperoxia exposure alters critical mitochondrial proteins related to cellular bioenergetics

We performed quantitative proteomics on ALI day 14 following hyperoxia exposure (ALI day 7 to 14) to define hyperoxia-sensitive alterations in the differentiated nTAEC proteome and to investigate how moderate hyperoxia alters pathways linking mitochondrial function, cellular metabolism and airway epithelial differentiation. Following MS acquisition of proteomics dataset, downstream analysis was performed with assessment of DEPs in response to hyperoxia. PCA was utilized to assess global proteomic differences between room air and hyperoxia-exposed term nTAECs. Biological replicates clustered distinctly by exposure condition, indicating a clear separation in global proteomic profiles between room air and hyperoxia groups (Fig 4A). For Volcano plot generation, we utilized a threshold of |log2FC| > 0.25 and FDR<0.05 to plot DEPs (Fig 4B). Of the 4895 unique proteins identified by DIA proteomics, hyperoxia exposure resulted in significant downregulation of 58 and upregulation of 73 DEPs, respectively. Among the top 20 downregulated DEPs (Fig 4C), hierarchical clustering and heatmap visualization revealed a striking suppression of multiple core mitochondrial respiratory Complex I components, including NDUFS1, NDUFS3, NDUFS4, NDUFS5, NDUFS7, NDUFS8, NDUFA5, NDUFA9, NDUFA12, NDUFB4, and NDUFV1, NDUFV2 (Fig. 4C). We further confirmed NDUFS1 protein expression in nTAECs with immunoblot (Fig S1) which demonstrated significant downregulation in the hyperoxia-exposed group (p=0.028). Complex I subunits are essential for the assembly and function of Complex I (48), the major entry point for electrons into the electron transport chain (ETC). Additional downregulated proteins included SDHB, a subunit of Complex II, and ACO1, a key tricarboxylic acid (TCA) cycle enzyme (49), indicating a broader disruption of mitochondrial bioenergetics.

**Figure 4.**
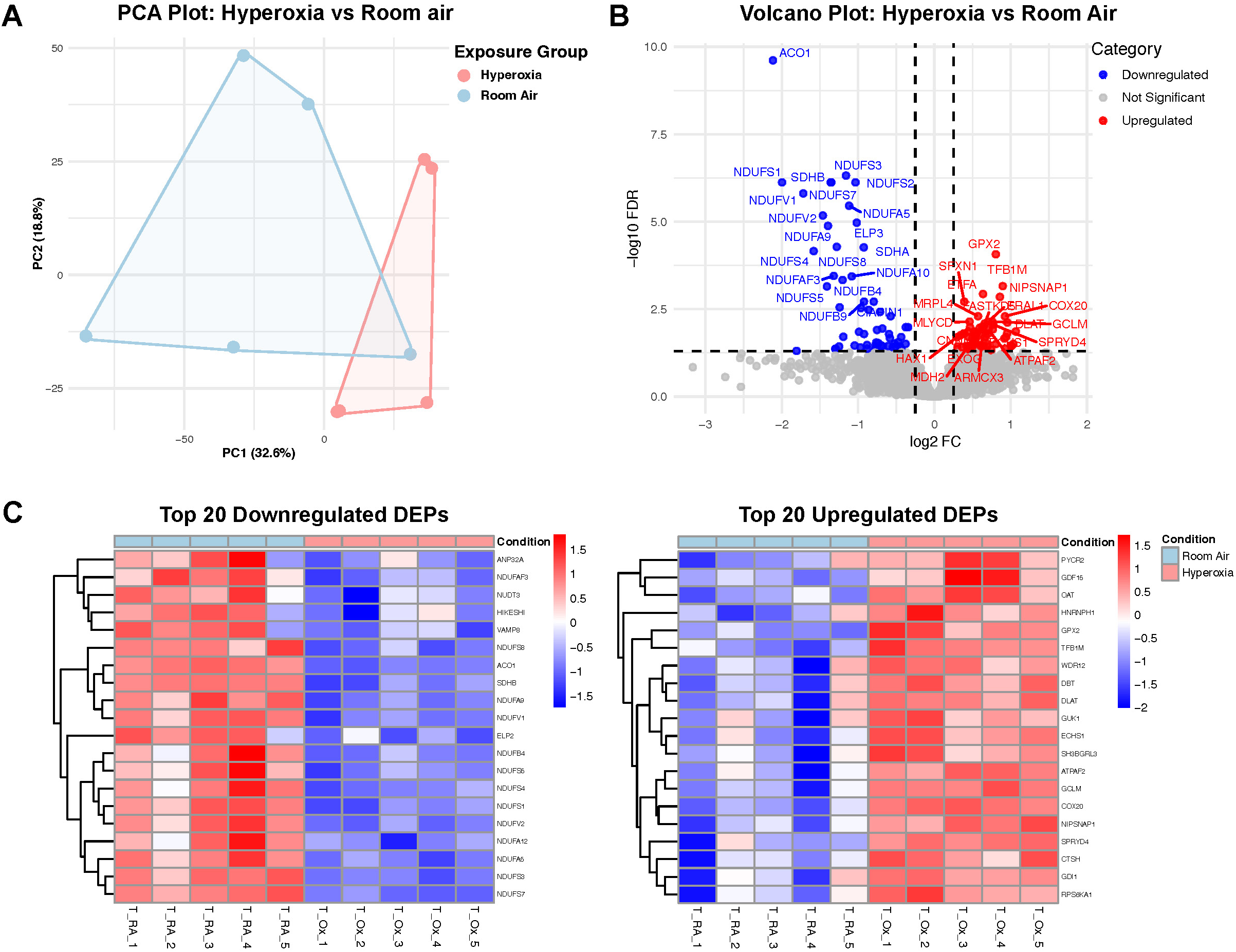
Hyperoxia disrupts mitochondrial function in neonatal tracheal airway epithelial cells through downregulation of critical electron transport chain (ETC) complexes. (A) Principal Component Analysis (PCA) was performed on log2-transformed, mean-imputed protein abundance values. The first two principal components (PC1 and PC2) explain 32.6% and 18.8% of the total variance, respectively. Each point represents an individual sample, colored by exposure group. Convex hulls highlight the clustering of biological replicates within each group, illustrating distinct global proteomic profiles between room air and hyperoxia-exposed nTAECs. (B) Volcano plot of differentially expressed proteins (DEPs) identified by Data Independent Acquisition (DIA) mass spectrometry in nTAECs on day 14 following 7 days of hyperoxia exposure (60% O₂, ALI day 7 to 14) compared to room air controls. Red and blue dots represent significantly upregulated and downregulated proteins, respectively (FDR ≤ 0.05, |log₂FC| > 0.25); gray dots represent proteins not meeting significance criteria. (C) Heatmaps showing top 20 significantly downregulated and upregulated DEPs ranked by FDR across room air and hyperoxia-exposed donor samples.

Conversely, the top 20 upregulated proteins in hyperoxia-exposed cells featured upregulation of GDF15, GPX2, GCLM, TFB1M, and ATPAF2 among others (Fig. 4C). These upregulated proteins include stress response mediators (GDF15, GPX2), redox-regulatory components (GCLM), and mitochondrial maintenance factors (TFB1M, ATPAF2), collectively suggesting a compensatory response to oxidative stress and impaired mitochondrial function under hyperoxic conditions.

To further interpret the biological significance of the DEPs in response to hyperoxia, we next performed comprehensive pathway enrichment analyses on the DEPs identified by quantitative proteomics. This integrative approach aimed to uncover converging molecular pathways related to mitochondrial function, metabolism, and hyperoxia induced epithelial injury responses relevant to the observed differentiation defects (41). GO BP analysis revealed that hyperoxia-exposed cells had significant enrichment of biological processes related to oxidative stress, reactive oxygen species (ROS) metabolism, and cellular redox homeostasis (Fig 5A). Reactome pathway enrichment showed similar findings, revealing suppression of respiratory ETC chain components (Fig 5B), especially genes involved in Complex I biogenesis and mitochondrial protein import (Fig S2), consistent with mitochondrial dysfunction observed in our Seahorse bioenergetics profiling. These pathways are known to support cellular differentiation processes through efficient mitochondrial ATP production and intermediate metabolism (50–52). We also applied IPA to evaluate both canonical signaling pathways and biological functions. Canonical pathway analysis demonstrated significant inhibition of oxidative phosphorylation and TCA cycle II, with concurrent activation of Sirtuin signaling, suggesting a compensatory stress-adaptive response (53) (Fig. 5C). IPA biological function prediction showed suppression of pathways involved in cell survival, protein metabolism, and cellular homeostasis, alongside activation of mitochondrial fragmentation, autophagy, and ROS synthesis (Fig. 5C).

**Figure 5.**
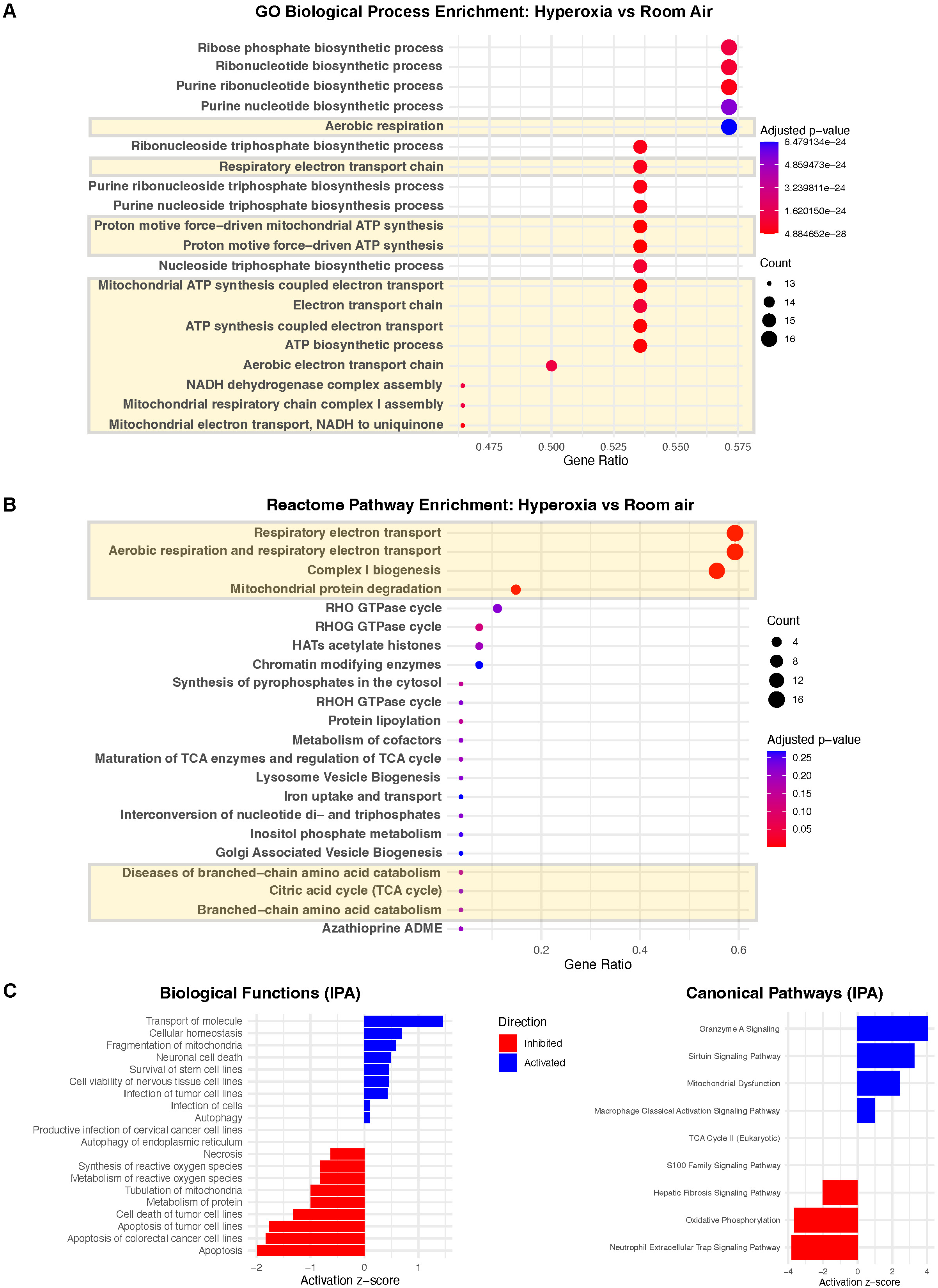
Pathway enrichment analyses of proteomic data identify disrupted oxidative phosphorylation and Complex I dysfunction as key features of hyperoxia-induced injury in neonatal airway epithelial cells. (A) Gene Ontology (GO) Biological Process enrichment analysis of all significant DEPs reveals significant overrepresentation of pathways involved in oxidative phosphorylation, NADH dehydrogenase activity, ATP biosynthesis, and electron transport. (B) Reactome pathway enrichment analysis of DEPs identifies significantly affected metabolic and mitochondrial processes, including respiratory electron transport, aerobic respiration, and Complex I biogenesis. Pathways are ranked by gene ratio and adjusted p-value. (C) Ingenuity Pathway Analysis (IPA) of DEPs highlight significantly altered biological functions and canonical pathways with activation z-scores indicating predicted directionality of pathway regulation.

### Gene Set Enrichment Analysis (GSEA) of the proteomic data links hyperoxia-induced mitochondrial dysfunction to metabolic reprogramming and epithelial differentiation pathways

To identify coordinated changes in biological pathways and cellular programs in response to hyperoxia exposure, we performed GSEA on the ranked proteomic dataset against the publicly available Reactome, WikiPathway and Hallmark pathway database. This approach enables the detection of subtle but biologically relevant alterations across functionally related proteins that may not meet conventional differential expression thresholds. The analysis focused on pathways related to mitochondrial function, metabolic regulation, and epithelial programs based on our prior hypothesis. Notably, Reactome ETC gene set revealed proteins involved in oxidative phosphorylation were significantly downregulated in the hyperoxia group, as evidenced by strong negative enrichment (NES = –3.32, FDR < 0.001) (Fig. 6A). Conversely, we observed significant positive enrichment for the Glucose Metabolism pathway (NES = +2.33, FDR = 0.027) in hyperoxia-exposed nTAECs (Fig. 6B). This included upregulation of HK2, ENO1, and GPI, indicative of a metabolic shift toward glycolysis (54) and consistent with a compensatory response to impaired oxidative phosphorylation and mitochondrial ATP production, complementing our finding from metabolic flux experiment in nTAECs in response to hyperoxia. Hallmark Fatty Acid Metabolism showed modest enrichment in the hyperoxia group (NES = +1.51, nominal p = 0.0724, FDR = 0.2919) (Fig. S3). Core enriched genes included several key enzymes involved in mitochondrial β-oxidation (55), including CPT1A, ACADM, ACADVL, ACAA1, HADHB, and ECHS1, suggesting a shift toward fatty acid catabolism as a compensatory mechanism to meet cellular energy demands under conditions of mitochondrial dysfunction. We also observed significant enrichment in amino acid metabolism pathway (NES = +2.06, nominal p = 0.0079, FDR = 0.1183) in the hyperoxia group (Fig S4). Proteins involved in the metabolism of branched-chain amino acids (56), glutamate (57), and cysteine (58) were particularly affected (e.g., BCAT2, GLS, GLUD1, GCDH, MPST), consistent with metabolic shift in response to hyperoxia.

**Figure 6.**
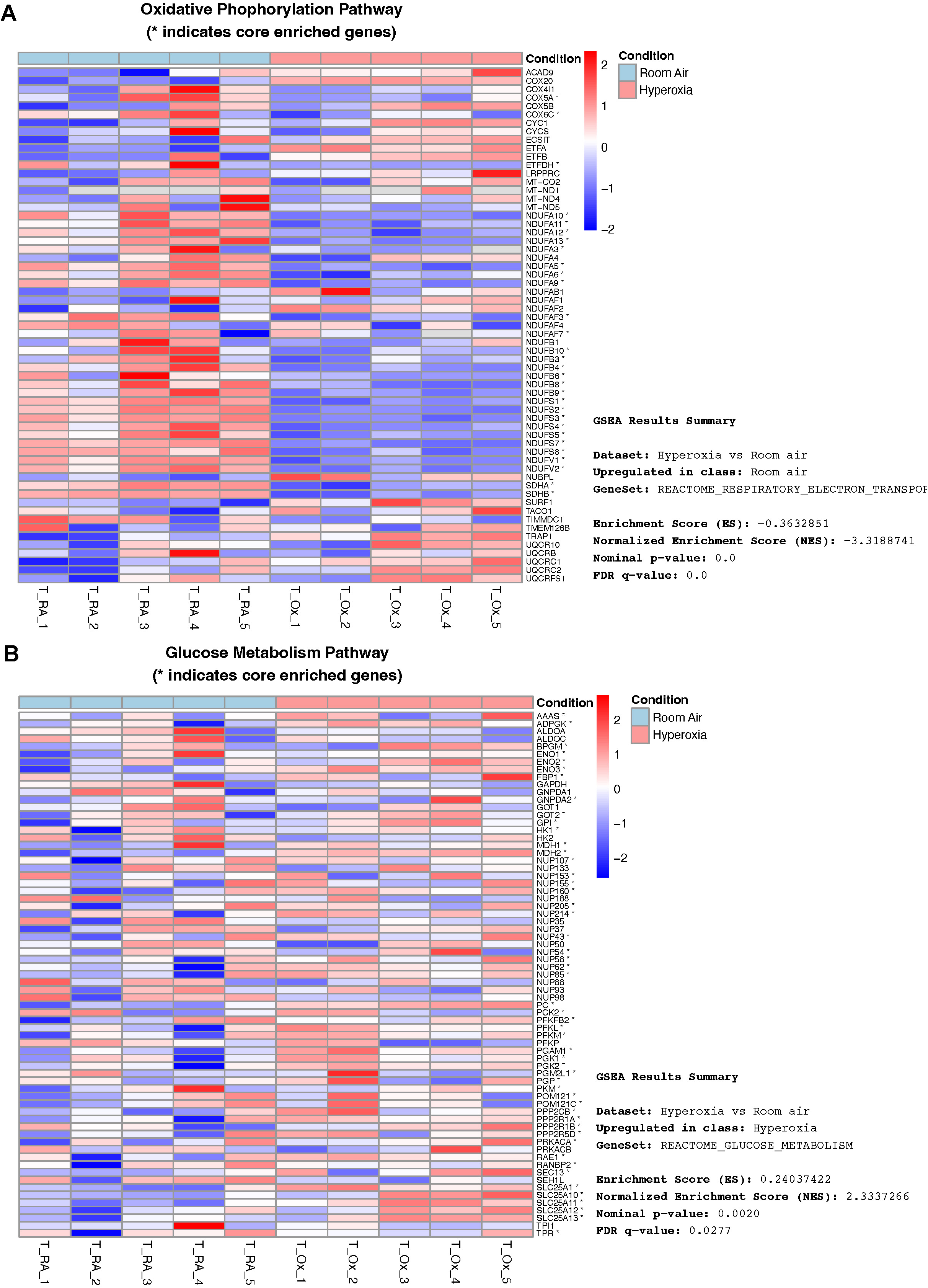
Gene Set Enrichment Analysis (GSEA) of proteomic dataset demonstrates that Hyperoxia downregulates oxidative phosphorylation and upregulates glucose metabolism in developing airway epithelial cells. (A) GSEA of DIA proteomic data reveals significant downregulation of proteins involved in the Reactome respiratory electron transport pathway in hyperoxia-exposed nTAECs on ALI day 14 (Normalized Enrichment Score [NES] = – 3.32, FDR q-value < 0.001). Heatmap shows expression of proteins in the Reactome respiratory electron transport gene set. Asterisks denote core enriched proteins contributing most to the enrichment score. (B) GSEA analysis identifies significant upregulation of the Reactome glucose metabolism pathway in response to hyperoxia (NES = 2.33, FDR q-value = 0.028). Heatmap displays protein expression across donor samples for this gene set, with core enriched proteins marked by asterisks.

Among the epithelial programs, Pre-Notch Expression and Processing pathway was positively enriched in the hyperoxia group (NES = +2.04, FDR = 0.139), suggesting activation of NOTCH signaling machinery, including upregulated expression of NOTCH2 and POGLUT1 (Fig S5). This is particularly noteworthy as NOTCH signaling has been implicated in regulating airway epithelial mucociliary differentiation (59). Finally, while the TGF-β signaling pathway gene set did not meet conventional significance thresholds (NES = 1.13, FDR q = 0.53), several core proteins including SERPINE1, RHOA, and THBS1 were upregulated in hyperoxia-exposed cells (Fig. S6) including the stress mitokine GDF15 (Fig 4C) which is an atypical member of the TGF-β superfamily (60) and has recently been shown to play important roles in neonatal hyperoxic lung injury (61, 62).

## DISCUSSION

To our knowledge, this is the first report integrating mitochondrial function analysis with DIA based quantitative proteomics in a human neonatal model of hyperoxia-exposed airway epithelium. This combined approach provides novel insights into how hyperoxia disrupts cellular bioenergetics and induces a shift in cellular metabolic pathways, influencing epithelial fate decisions during early lung development. Our findings build upon previous studies linking hyperoxia to neonatal airway epithelial injury (29, 63), including animal models that demonstrate altered club-to-ciliated cell ratios in response to oxidative stress (64). Notably, no prior work has utilized neonatal tracheal aspirate-derived airway epithelial cells in an organotypic culture to study epithelial differentiation programs under clinically relevant environmental oxygen exposure conditions.

We demonstrate that moderate hyperoxia (60% O_2_) exposure during mid-phase of mucociliary differentiation impairs terminal differentiation of neonatal patient-derived nTAECs. In our model, hyperoxia stunts epithelial maturation resulting in persistence of a progenitor-like state (e.g., basal and club cells) and decreased terminal differentiation into goblet and ciliated cells, without significant impact in cell monolayer integrity. These features are reminiscent of the early canalicular phase of the developing lung where the airway epithelium is predominantly populated by basal and club cells. Seahorse metabolic flux analysis of nTAECs demonstrates that moderate hyperoxia reprograms the metabolic phenotype of nTAECs, characterized by reduced oxidative phosphorylation and enhanced glycolysis, indicative of mitochondrial bioenergetic dysfunction. Given that cellular differentiation, especially terminal epithelial maturation including ciliary development and mucus production, are energy intensive processes (50–52), we speculate that the hyperoxia-induced cellular metabolic shift impedes terminal differentiation in nTAECs. To uncover molecular mechanisms that underlie hyperoxia-induced differentiation defect and bioenergetic dysfunction, we performed DIA proteomic study following hyperoxia exposure. Quantitative proteomic profiling revealed downregulation of mitochondrial ETC Complex I and II subunits (e.g., NDUFS1, NDUFV1 and SDHB among others) and TCA cycle enzymes in response to hyperoxia. Complex I serves as the major entry point for electrons into the ETC and is essential for oxidative phosphorylation (48). Its dysfunction impairs mitochondrial ATP generation and promotes ROS accumulation (49), both of which can interfere with signaling pathways guiding epithelial differentiation. This was accompanied by an increase in stress-responsive cytokines (GDF15, GPX2, GCLM, TFB1M, and ATPAF2 among others) in the hyperoxia group. These proteins are involved in cellular stress responses, redox regulation, and compensatory mitochondrial remodeling (46). For example, GDF15 is a known mitochondrial stress-responsive cytokine (60), while GPX2 (65) and GCLM (66) are involved in glutathione metabolism and oxidative stress mitigation. Upregulation of TFB1M (67), a mitochondrial transcription factor, and ATPAF2 (68), a Complex V assembly factor, suggests activation of adaptive responses to preserve mitochondrial integrity under oxidative conditions.

Cellular bioenergetic function has previously been linked to later development of BPD. A 2022 study (69) compared the mitochondrial function of umbilical cord-derived mesenchymal stem cells from extremely preterm infants who subsequently developed severe BPD to those from infants who did not or developed only mild disease. The cells from infants who went on to develop severe BPD had lower OCR and higher ECAR indicating a metabolic shift reminiscent of that observed in hyperoxia-exposed nTAECs in our model. Another study with nasopharyngeal suction aspirate-derived airway epithelial cells from term and moderately preterm infants showed that preterm cells had a lower OCR at baseline and failed to reach maximal respiratory capacity when challenged as compared to term cells (28). Intriguingly, preterm neonatal airway epithelial cells had significantly lower gene expression of NDUFA6, a critical Complex I subunit, which might indicate a potentially diminished Complex I function at baseline in preterm airway epithelial cells. Further supporting this, Teape et al. (29) demonstrated that hyperoxia exposure led to a time-dependent reduction in cilia length and ciliated surface area in adult-derived airway epithelial cells. In a complementary neonatal hyperoxia mouse model, they observed reduced expression of ACTUB in the airway following hyperoxia (>95% O_2_) exposure. Furthermore, their proteomic and metabolomic data also revealed a significant downregulation of multiple Complex I subunits (including NDUFA7 and NDUFS4), aligning with our findings of impaired mitochondrial function as evidenced in their model by a decreased ATP/ADP ratio. In our model, we demonstrate that the mid-phase of airway epithelial differentiation represents a critical window of vulnerability, during which hyperoxia induces mitochondrial dysfunction and a metabolic shift, coinciding with impaired epithelial differentiation. Notably, this vulnerability was evident even under clinically relevant levels of hyperoxia (60% O₂), which was sufficient to replicate the differentiation defects observed in prior studies utilizing more severe hyperoxia (>95% O₂) exposure strategies. This suggests that even moderate hyperoxia exposure during key developmental stages can disrupt mitochondrial function and impair airway epithelial maturation.

Downstream pathway enrichment analyses of our proteomics data provide further mechanistic insight into hyperoxia-sensitive molecular programs related to mitochondrial function and airway epithelial maturation. GO and Reactome enrichment analyses revealed significant activation of oxidative stress-related processes, including ROS metabolism and redox homeostasis, while concurrently demonstrating suppression of respiratory ETC components, particularly those involved in Complex I biogenesis and mitochondrial protein import. These findings align with our observed suppression of mitochondrial ATP production in seahorse analysis and suggest that impaired mitochondrial energetics may compromise the cellular metabolic demand required for terminal epithelial differentiation. Complementary IPA further substantiates these observations, revealing broad inhibition of canonical bioenergetic pathways such as oxidative phosphorylation and the TCA cycle, coupled with compensatory activation of sirtuin signaling (53), which is a known adaptive response to cellular stress. IPA biological function prediction highlighted suppression of essential processes like protein metabolism, cell survival, and homeostasis, while simultaneously predicting activation of mitochondrial fragmentation and autophagy of the endoplasmic reticulum, which implicates escalating subcellular organelle stress (70) as a key effector mechanism of epithelial injury. Prior studies have demonstrated that autophagy plays a critical role in airway epithelial injury by maintaining and priming the stem/progenitor cell pool necessary for effective tissue repair (71). In addition to autophagy, glucose metabolism was also identified as a key factor in preserving the epithelial stem cell niche. Taken together, our findings support a model in which hyperoxia-induced mitochondrial dysfunction and metabolic reprogramming activate epithelial injury-response programs that sustain a progenitor-like state. While this may be protective in the context of injury repair, it may also hinder terminal differentiation, ultimately impairing epithelial maturation.

GSEA of our proteomic dataset highlights a global shift in the metabolic programming of nTAECs following hyperoxia exposure. Suppression of the ETC and oxidative phosphorylation pathways, particularly through downregulation of Complex I subunits, was accompanied by enrichment of glycolysis and glucose metabolism pathways, suggesting a metabolic shift toward glycolytic ATP production, directly correlating with the hyperoxia-induced metabolic alteration observed during metabolic flux analysis of nTAECs. This metabolic shift was further supported by modest enrichment of fatty acid β-oxidation and branched-chain amino acid metabolism, indicating compensatory use of alternative energy substrates. Intriguingly, MPST, a key mitochondrial enzyme in the transsulfuration pathway (58, 72), was downregulated in hyperoxia. MPST is known to contribute towards mitochondrial redox balance through a specialized post-translational modification called protein persulfidation, which has been shown to protect mitochondrial Complex I from oxidative damage in cardiac and adipose tissues (73). MPST is also involved in mitochondrial protein import and preserving cellular bioenergetic function (74, 75). Its reduction suggests compromised mitochondrial sulfide signaling, further sensitizing cells to oxidative stress. In parallel with these metabolic changes, our GSEA data reveal dysregulation of key epithelial differentiation pathways. The Pre-Notch Expression and Processing pathway was positively enriched in hyperoxia-exposed cells, consistent with activation of NOTCH signaling, a master regulator of airway epithelial lineage balance (59). NOTCH signaling activation is known to suppress ciliated cell fate while promoting secretory cell differentiation, particularly club and goblet cells. While our observation of reduced ciliated cells in response to hyperoxia aligns with this paradigm, the concurrent decrease in goblet cells is unexpected. We speculate that hyperoxia-induced mitochondrial dysfunction and energy deficits may impair the terminal differentiation of goblet cells, despite upstream NOTCH activation. Additionally, modest enrichment of the TGF-β signaling pathway and upregulation of stress-responsive effectors such as GDF15, SERPINE1, and THBS1 suggest that oxidative stress may activate autocrine or paracrine TGF-β signaling in this context. TGFβ plays essential roles during normal lung development, but its dysregulation is strongly associated with adverse respiratory outcomes in preterm infants (76–79). TGFβ dysregulation plays an important role in airway remodeling in both BPD (80, 81) and COPD (82). The effects of TGF-β signaling on airway epithelial differentiation are context-dependent. A prior study demonstrated that even under pro-goblet conditions, such as stimulation with interleukin-13 (IL-13), active SMAD signaling, a canonical downstream pathway of TGF-β, can suppress goblet cell differentiation This provides a plausible mechanistic explanation for our observed phenotype in that hyperoxia-induced oxidative stress might activate TGF-β/SMAD signaling, which in turn may block the terminal differentiation of secretory cells, including goblet cells, despite concurrent NOTCH-driven lineage priming. These findings also highlight that hyperoxia-induced mitochondrial dysfunction and metabolic shift may exert cell-type specific effects on epithelial programs, disrupting normal lineage progression through distinct but overlapping mechanisms.

The strength of our study lies in the innovative use of neonatal patient-sample derived nTAECs as a translational model (31) along with clinically relevant oxygen exposure strategy. In contrast to prior studies that have primarily utilized adult-derived epithelial cells, our approach addresses the unique developmental vulnerability of the neonatal airway epithelium, which is central to diseases such as BPD (10, 15). Additionally, the innovative use of DIA quantitative proteomics is a major strength of our study. This platform enabled high-resolution mapping of proteomic changes in response to hyperoxia, allowing us to uncover critical alterations in mitochondrial pathways and metabolic signaling and has been instrumental in generating mechanistic hypotheses that link bioenergetic dysfunction to impaired epithelial differentiation. Our study is not without limitations. We have used nTAECs isolated from term infants as opposed to infants with established BPD. We intentionally selected this study population to align with our central objective of investigating the impact of hyperoxia on neonatal airway epithelial development. Including samples from infants with complicated clinical courses, such as those who develop BPD, could introduce significant donor-specific variability, making it difficult to isolate and interpret the specific effects of hyperoxia. To simulate the active epithelial development in extremely preterm infants, our model initiates hyperoxic exposure during the mid-phase of airway epithelial differentiation, rather than after full maturation as in prior models(29). This approach more accurately reflects the *in vivo* timing of injury during neonatal lung development. We performed cellular bioenergetic profiling using undifferentiated basal cells due to technical limitations associated with conducting the assay on transwell inserts especially after the formation of a mature epithelial monolayer. However, we plan to incorporate previously published methods in future studies to directly assess cellular bioenergetics in transwell inserts (83). While DIA proteomics is a robust and unbiased approach, it may fail to detect low-abundant proteins which could be biologically significant (85). Additionally, the use of whole-cell lysates for proteomic analysis limits the ability to resolve cell type-specific protein expression and subcellular localization. This approach may obscure important spatial and functional differences between distinct epithelial cell populations, such as basal, ciliated, and secretory cells, which could be critical for interpreting the effects of hyperoxia on airway epithelial differentiation and function. Further, our study was limited by a relatively small sample size and was not adequately powered to detect potential sex-specific differences in the epithelial response to hyperoxia. Finally, these findings are based on an *in vitro* model and will require validation through *in vivo* studies using mouse models, which are currently underway.

Our future studies will explore whether restoring or prophylactically boosting mitochondrial function through pharmacologic or genetic interventions can rescue the abnormal epithelial phenotype. Additionally, we intend to isolate nTAECS from neonates with established BPD and compare the epithelial phenotype with our current model of hyperoxia exposed nTAECs. Finally, the inherent heterogeneity of the human airway epithelium means that hyperoxia-induced mitochondrial dysfunction likely exerts cell-type specific effects on epithelial differentiation programs. Therefore, integration of single-cell approaches, as demonstrated in recent robust studies (86, 87) of neonatal lung injury, will be instrumental in delineating lineage-specific disruptions and identifying vulnerable subpopulations within the developing airway in response to hyperoxic injury.

## CONCLUSION

In summary, our study identifies mitochondrial bioenergetic dysfunction as a key mediator of hyperoxia-induced impairment in neonatal airway epithelial development. Complementary *in vivo* studies are warranted to further model these effects, and targeting mitochondrial health may represent a novel therapeutic strategy to mitigate long-term airway disease in survivors of hyperoxia-induced BPD.

## SUPPLEMENTAL MATERIALS

Supplemental Table S1 and S2, Supplemental Method and Supplemental Figs. S1-S6. Supplemental materials are uploaded in FigShare and can be accessed with this private link: https://figshare.com/s/bab22d0cd5753e14c6d6

## ACKNOWLEDGEMENTS

We would like to acknowledge Ms. Erin Bohon and Aprill Shockley, Neonatal Research Nurses at Oklahoma Children’s Hospital, for their substantial support regarding screening and consenting parents, collection of neonatal samples and transport to the lab. We would like to thank the IDeA National Resource for Quantitative Proteomics for generous help with proteomic experiments and subsequent data analysis. Graphical images were created with Biorender. Finally, we would like to acknowledge utilization of ChatGPT (OpenAI, 2025 version) to assist with English language editing and improving clarity in sections of the manuscript. All AI-generated content was reviewed and verified by the authors for accuracy and integrity.

## AUTHOR CONTRIBUTIONS

A.G., T.E.T and A.R. conceived and designed the experiments; A.G., C.C. and A.R.N performed experiments; A.G., H.Z. and A.M. analyzed data; A.G., L.K.R, M.S.W, T.E.T and A.R. interpreted results; A.G. prepared figures and drafted the manuscripts; A.G., L.K.R, M.S.W, Y.S.P, T.E.T and A.R. edited and revised manuscript; All the authors read and approved the final version of the manuscript.

## DATA AVAILABILITY

The mass spectrometry proteomics data have been deposited to the ProteomeXchange Consortium via the PRIDE partner repository with the dataset identifier PXD064170 and 10.6019/PXD064170. The corresponding author will provide original data upon a reasonable request.

