## Supplemental Table 1 for "Quantitative Proteomics Links Mitochondrial Dysfunction to Metabolic Changes and Epithelial Differentiation Defects in Hyperoxia-Exposed Neonatal Airway Cells"

**Table S1:** Clinical characteristics of infants

| **ID** | **Gestational age at birth** | **Sex** | **Clinical diagnosis at NICU admission** |
| --- | --- | --- | --- |
| 1 | 37 weeks 0 days | M | Hypoxic-Ischemic Encephalopathy |
| 2 | 38 weeks 0 days | M | Congenital Heart Disease |
| 3 | 37 weeks 2 days | M | GI pathology |
| 4 | 37 weeks 0 days | F | Hypoxic-Ischemic Encephalopathy |
| 5 | 38 weeks 3 days | M | GI pathology |
