## Supplemental Table 2 for "Quantitative Proteomics Links Mitochondrial Dysfunction to Metabolic Changes and Epithelial Differentiation Defects in Hyperoxia-Exposed Neonatal Airway Cells"

**Table S2:** Key proteins identified throughout the study including quantitative proteomic analyses and their associated pathways

| **Gene Symbol** | **Protein Name** | **Pathway** |
| --- | --- | --- |
| ACAA1 | 3-Ketoacyl-CoA Thiolase, Peroxisomal | Fatty Acid β-Oxidation |
| ACADM | Acyl-CoA Dehydrogenase Medium Chain | Fatty Acid β-Oxidation |
| ACADVL | Acyl-CoA Dehydrogenase Very Long Chain | Fatty Acid β-Oxidation |
| ACO1 | Aconitase 1 | TCA Cycle / Electron Transport Chain |
| ACTUB | Acetylated Tubulin | Epithelial Cell Markers |
| ATPAF2 | ATP Synthase Mitochondrial F1 Complex Assembly Factor 2 | Mitochondrial Stress and Biogenesis |
| BCAT2 | Branched Chain Amino Acid Transaminase 2 | Amino Acid Metabolism |
| CPT1A | Carnitine Palmitoyltransferase 1A | Fatty Acid β-Oxidation |
| ECHS1 | Enoyl-CoA Hydratase 1 | Fatty Acid β-Oxidation |
| ENO1 | Alpha-Enolase | Glycolysis |
| FOXJ1 | Forkhead Box J1 | Epithelial Cell Markers |
| GCDH | Glutaryl-CoA Dehydrogenase | Amino Acid Metabolism |
| GCLM | Glutamate-Cysteine Ligase Modifier Subunit | Mitochondrial Stress and Biogenesis |
| GDF15 | Growth Differentiation Factor 15 | Mitochondrial Stress and Biogenesis/TGF-β Pathway |
| GLS | Glutaminase | Amino Acid Metabolism |
| GLUD1 | Glutamate Dehydrogenase 1 | Amino Acid Metabolism |
| GPI | Glucose-6-Phosphate Isomerase | Glycolysis |
| GPX2 | Glutathione Peroxidase 2 | Mitochondrial Stress and Biogenesis |
| HADHB | Hydroxyacyl-CoA Dehydrogenase Trifunctional Multienzyme Complex Subunit Beta | Fatty Acid β-Oxidation |
| HK2 | Hexokinase 2 | Glycolysis |
| MUC5AC | Mucin 5AC | Epithelial Cell Markers |
| NDUFA12 | NADH:Ubiquinone Oxidoreductase Subunit A12 | Mitochondrial Complex I (ETC) |
| NDUFA5 | NADH:Ubiquinone Oxidoreductase Subunit A5 | Mitochondrial Complex I (ETC) |
| NDUFA9 | NADH:Ubiquinone Oxidoreductase Subunit A9 | Mitochondrial Complex I (ETC) |
| NDUFB4 | NADH:Ubiquinone Oxidoreductase Subunit B4 | Mitochondrial Complex I (ETC) |
| NDUFS1 | NADH:Ubiquinone Oxidoreductase Core Subunit S1 | Mitochondrial Complex I (ETC) |
| NDUFS3 | NADH:Ubiquinone Oxidoreductase Core Subunit S3 | Mitochondrial Complex I (ETC) |
| NDUFS4 | NADH:Ubiquinone Oxidoreductase Subunit S4 | Mitochondrial Complex I (ETC) |
| NDUFS5 | NADH:Ubiquinone Oxidoreductase Subunit S5 | Mitochondrial Complex I (ETC) |
| NDUFS7 | NADH:Ubiquinone Oxidoreductase Subunit S7 | Mitochondrial Complex I (ETC) |
| NDUFS8 | NADH:Ubiquinone Oxidoreductase Subunit S8 | Mitochondrial Complex I (ETC) |
| NDUFV1 | NADH:Ubiquinone Oxidoreductase Flavoprotein 1 | Mitochondrial Complex I (ETC) |
| NDUFV2 | NADH:Ubiquinone Oxidoreductase Flavoprotein 2 | Mitochondrial Complex I (ETC) |
| NOTCH2 | Neurogenic locus notch homolog protein 2 | NOTCH Signaling |
| POGLUT1 | Protein O-glucosyltransferase 1 | NOTCH Signaling |
| RHOA | Ras Homolog Family Member A | TGF-β Pathway / Epithelial Remodeling |
| SCGB1A1 | Secretoglobin Family 1A Member 1 (Club Cell Secretory Protein) | Epithelial Cell Markers |
| SDHB | Succinate Dehydrogenase Complex Iron Sulfur Subunit B | TCA Cycle / Electron Transport Chain |
| SERPINE1 | Serpin Family E Member 1 | TGF-β Pathway / Epithelial Remodeling |
| TFB1M | Transcription Factor B1, Mitochondrial | Mitochondrial Stress and Biogenesis |
| THBS1 | Thrombospondin 1 | TGF-β Pathway / Epithelial Remodeling |
| P63-⍺ | Tumor Protein P63 Alpha Isoform | Epithelial Cell Markers |
