## Supplemental Method and Figures for "Quantitative Proteomics Links Mitochondrial Dysfunction to Metabolic Changes and Epithelial Differentiation Defects in Hyperoxia-Exposed Neonatal Airway Cells"

**Western Blotting**

Total protein was isolated by lysing cells directly within the ALI culture wells using 100 µL of RIPA buffer (Cat# R0278, Sigma-Aldrich) supplemented with a cocktail of protease inhibitors (1X, Cat# 88666, Thermo Scientific), phosphatase inhibitors (1X, Cat# 1862495, Thermo Scientific), Laemmli sample buffer (1X, Cat# 1610747, Bio-Rad), and 2.5% 2-mercaptoethanol (Cat# 21985023, Thermo Scientific). Protein lysates were resolved using 4-20% SDS-PAGE gels and subsequently transferred onto nitrocellulose membranes via semi-dry transfer. Membranes were blocked for 1 hour at room temperature in 5% non-fat dry milk in TBST and incubated overnight at 4°C with the primary antibodies NDUFS1 (12444-1-AP, 1:1000, Proteintech). The following day, membranes were washed and incubated for 1 hour at room temperature with horseradish peroxidase (HRP)-conjugated secondary antibodies: goat anti-mouse HRP (1:2000, Cat# 1031-05, Southern Biotech) or goat anti-rabbit HRP (1:2000, Cat# 4030-05, Southern Biotech). Detection was performed using enhanced chemiluminescence (ECL), and protein bands were imaged with the ChemiDoc system. Band intensities were quantified by densitometry using Image Studio software (version 5.2.1, LI-COR, Lincoln, NE). All target protein levels were normalized to GAPDH (5174S, 1:5000, Cell Signaling Technology) as a loading control.

**SUPPLEMENTARY FIGURES**:

**Figure S1.** Immunoblot for NDUFS1, a mitochondrial complex I subunit critical for maintaining mitochondrial bioenergetic function on ALI day 14 following hyperoxia exposure of nTAECs. Data points represent protein expression normalized to room air control for each donor (n = 4 donor cells). GAPDH was used as endogenous control. Statistical analysis was performed utilizing Mann-Whitney U test. Numbers (1 to 4) represent corresponding donors in room air and hyperoxia group.

**Figure S2:** (A) Gene Set Enrichment Analysis (GSEA) of DIA proteomic data reveals significant downregulation of proteins involved in Complex I biogenesis in hyperoxia-exposed nTAECs on ALI day 14 (Normalized Enrichment Score [NES] = –3.84, FDR q-value < 0.001). Heatmap shows expression of proteins in the Reactome Complex I gene set. Asterisks denote core enriched proteins contributing most to the enrichment score. (B) GSEA analysis identifies significant upregulation of Reactome mitochondrial protein import pathway in response to hyperoxia (NES = 2.92, FDR q-value = 0.001). Heatmap displays protein expression across donor samples for this gene set, with core enriched proteins marked by asterisks.

**Figure S3:** Gene Set Enrichment Analysis (GSEA) of DIA proteomic data reveals enrichment of the Hallmark fatty acid metabolism pathway in hyperoxia-exposed nTAECs on ALI day 14 (Normalized Enrichment Score [NES] = 1.51, nominal *p* = 0.072, FDR q-value = 0.292). Heatmap displays protein expression of genes in the fatty acid metabolism pathway across donor samples. Asterisks denote core enriched proteins contributing most to the enrichment score. Proteins span key regulators of mitochondrial β-oxidation, including CPT1A, ACADM, ACADVL, HADHB, and ECHS1, supporting increased lipid catabolism as a metabolic adaptation to hyperoxic stress.

**Figure S4:** Gene Set Enrichment Analysis (GSEA) of DIA proteomic data identifies significant enrichment of the WikiPathway amino acid metabolism pathway in hyperoxia-exposed nTAECs on ALI day 14 (Normalized Enrichment Score [NES] = 2.06, nominal *p* = 0.0079, FDR q-value = 0.1183). Heatmap displays protein expression of genes in the amino acid metabolism pathway across donor samples, with asterisks indicating core enriched proteins contributing most to the enrichment score.

**Figure S5:** Gene Set Enrichment Analysis (GSEA) of DIA proteomic data shows significant enrichment of the Reactome Pre-Notch Expression and Processing pathway in hyperoxia-exposed nTAECs on ALI day 14 (Normalized Enrichment Score [NES] = 2.04, nominal *p* = 0.010, FDR q-value = 0.139). Heatmap displays protein expression of pathway members across donor samples, with asterisks indicating core enriched proteins contributing most to the enrichment score.

**Figure S6:** (A) Gene Set Enrichment Analysis (GSEA) of DIA proteomic data shows a trend toward enrichment of the Hallmark TGF-β signaling pathway in hyperoxia-exposed nTAECs on ALI day 14 (Normalized Enrichment Score [NES] = 1.13, nominal *p* = 0.285, FDR q-value = 0.528). (B) Heatmap displays protein expression across donor samples for the TGF-β signaling pathway. Asterisks denote core enriched proteins contributing most to the enrichment score. Although the gene set did not meet conventional significance thresholds, several stress-activated or Epithelial Mesenchymal Transition (EMT)-associated effectors such as SERPINE1, THBS1, and RHOA were upregulated in response to hyperoxia, suggesting enhanced TGF-β–related signaling and epithelial remodeling under oxidative stress.

**FIGURE S1**

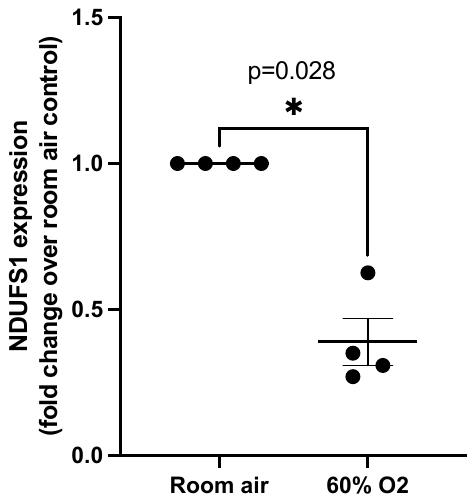

**FIGURE S2**

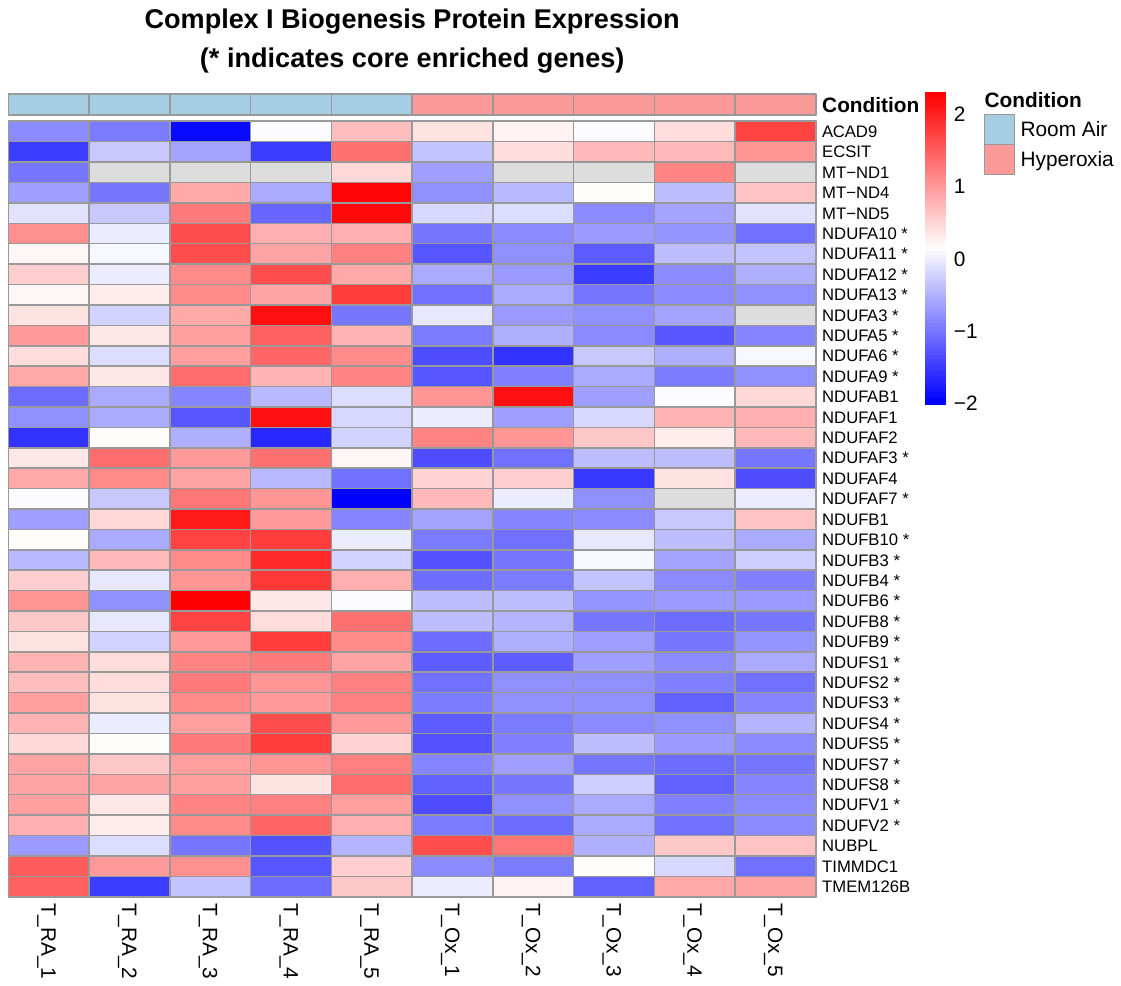

**(A)**

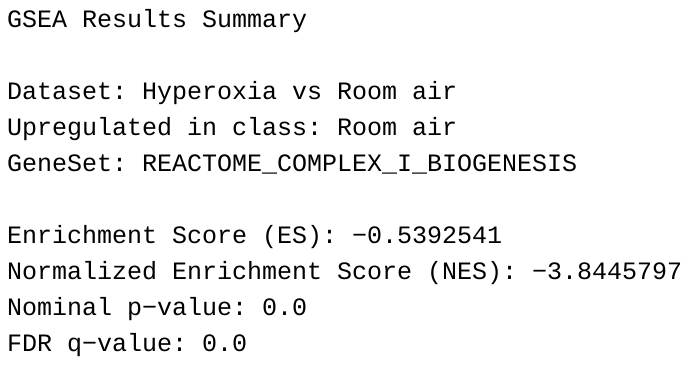

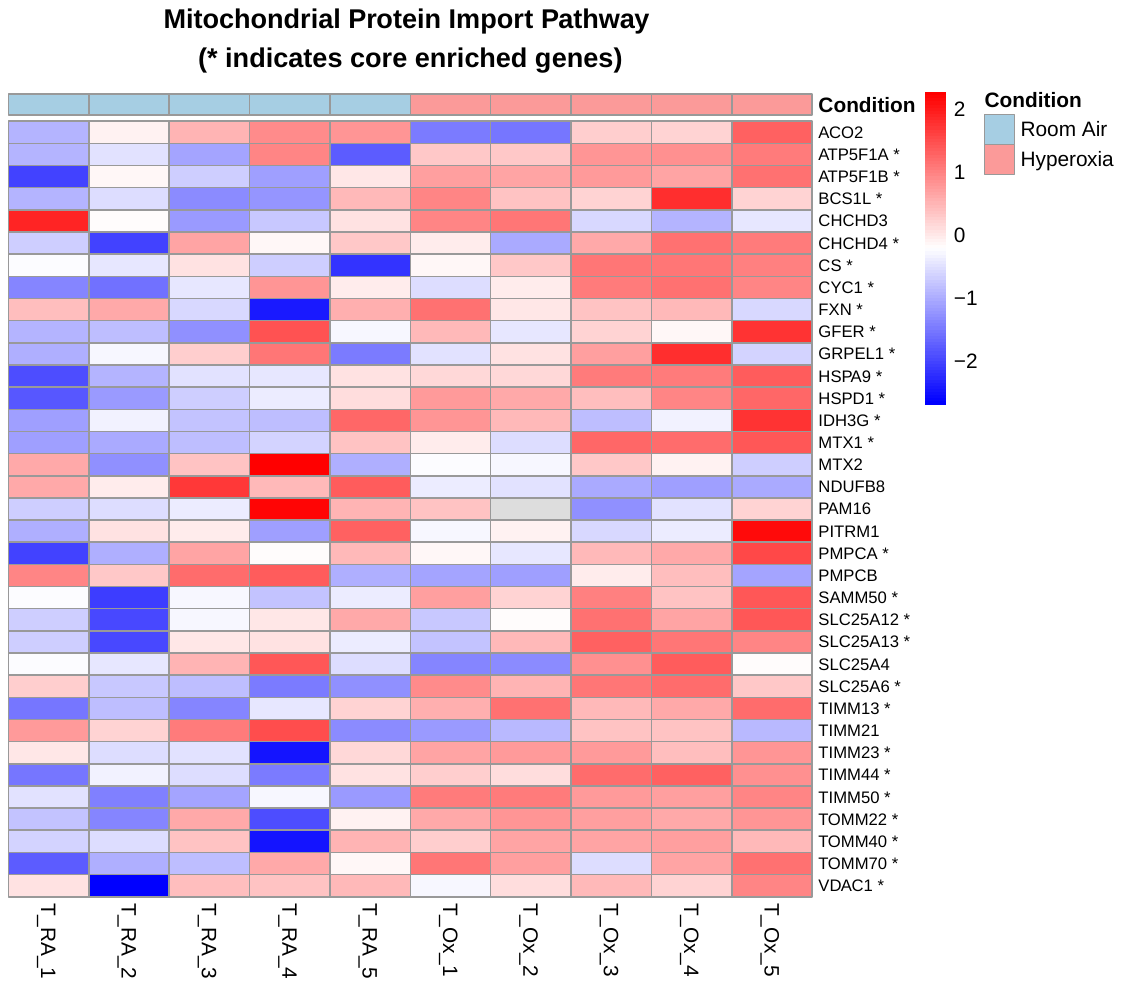

**(B)**

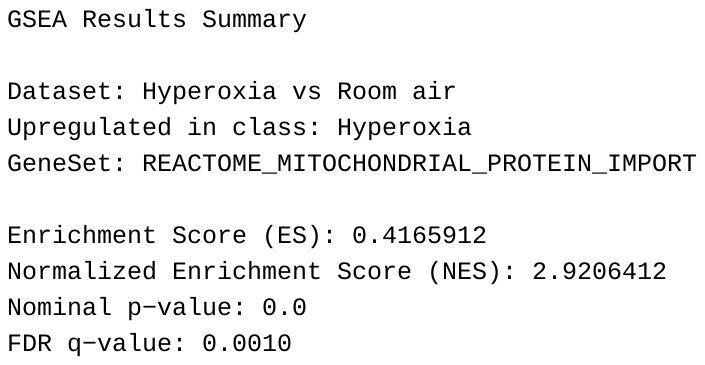

**FIGURE S3**

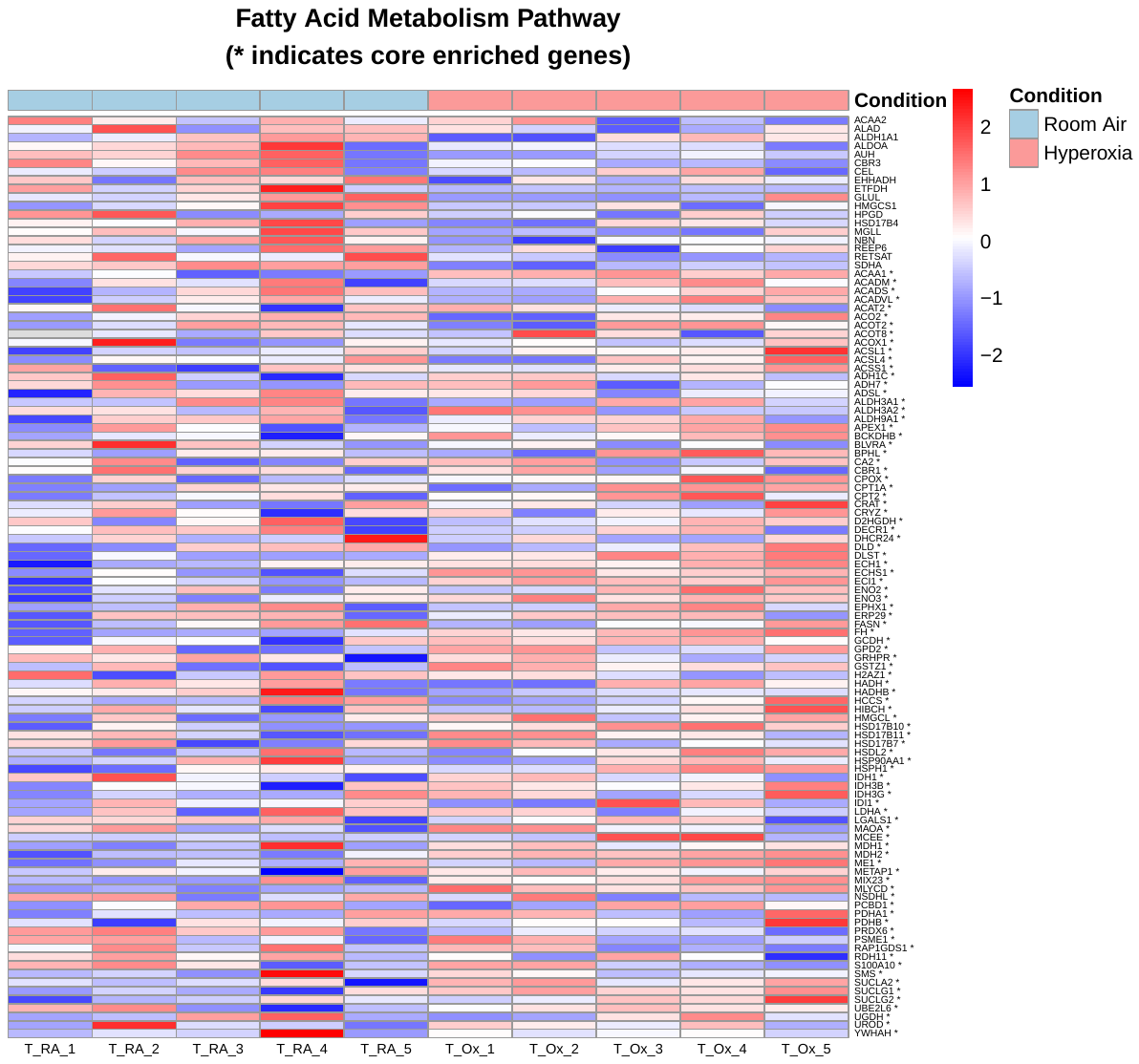

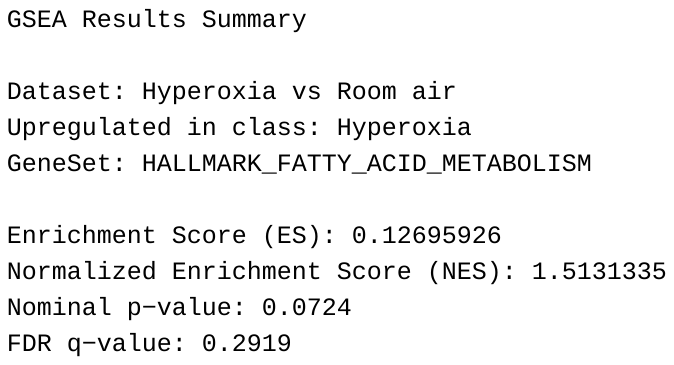

**FIGURE S4**

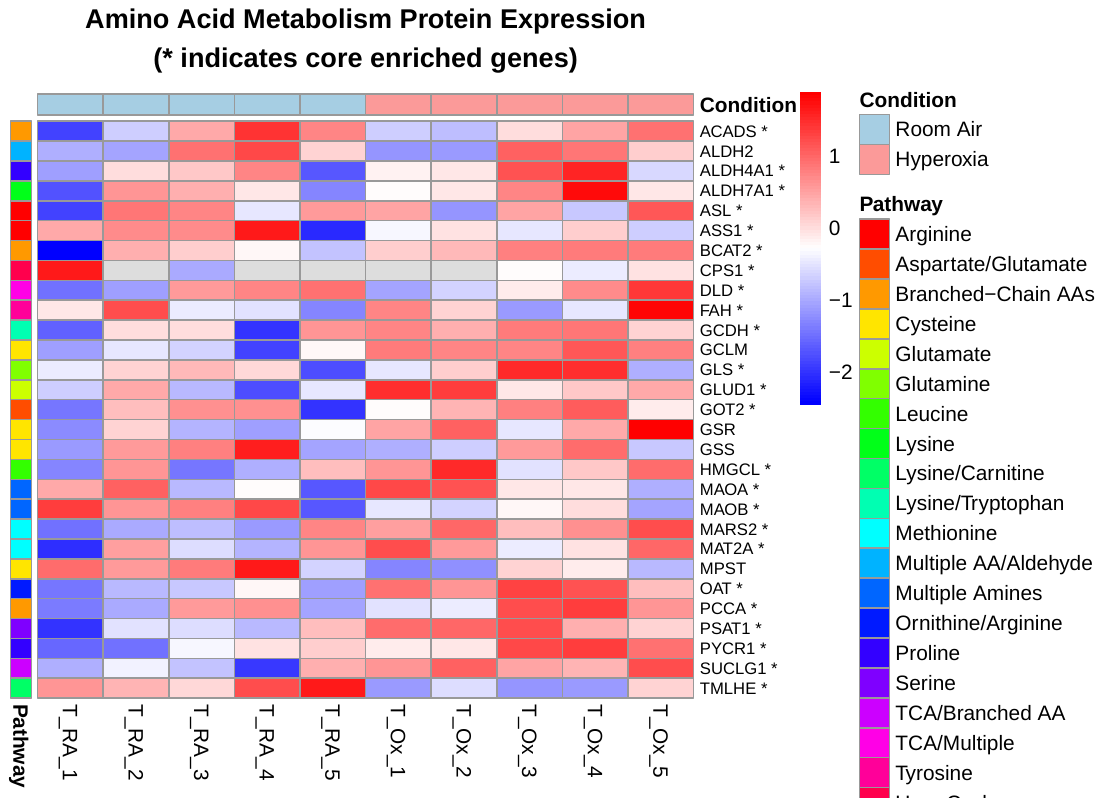

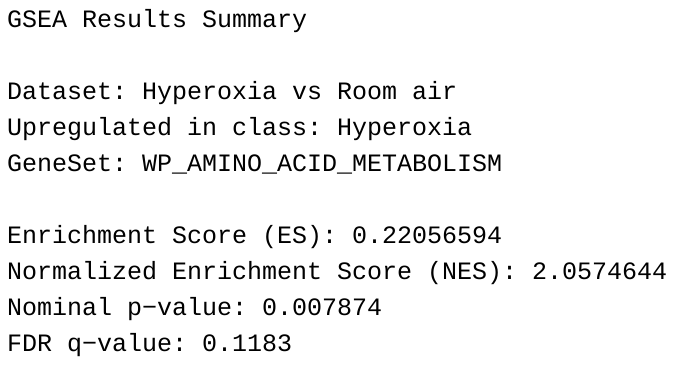

**FIGURE S5**

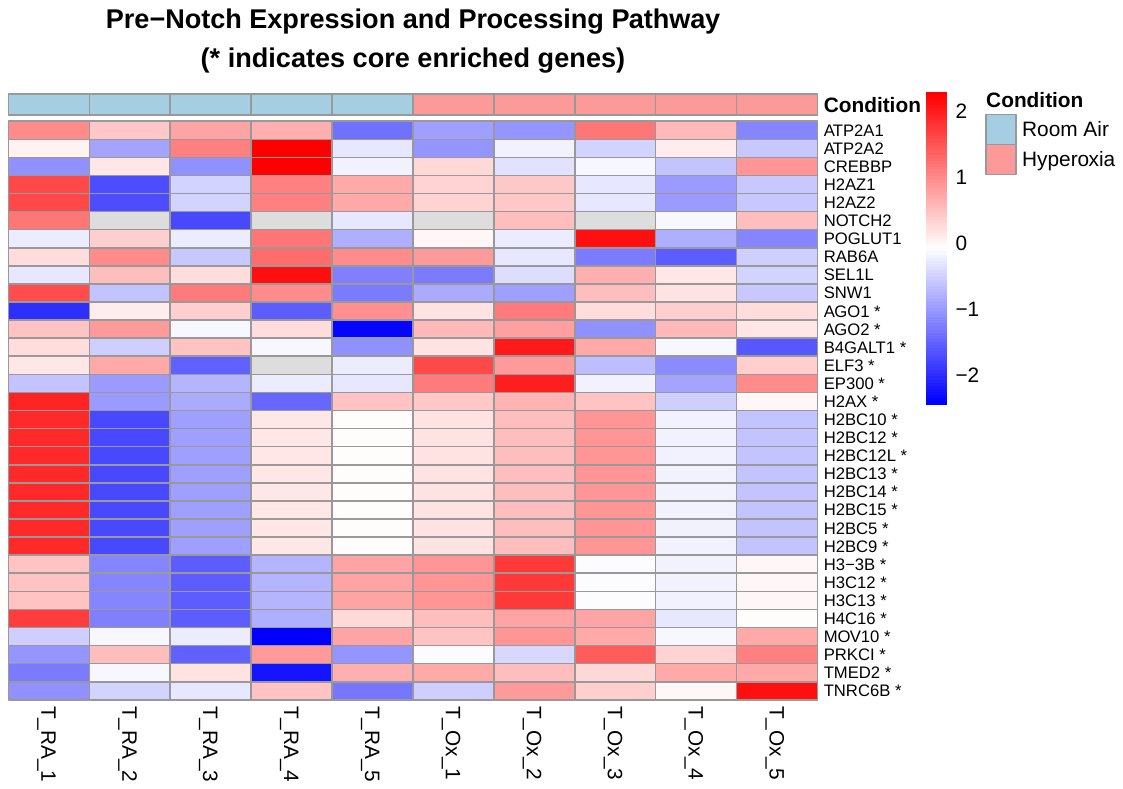

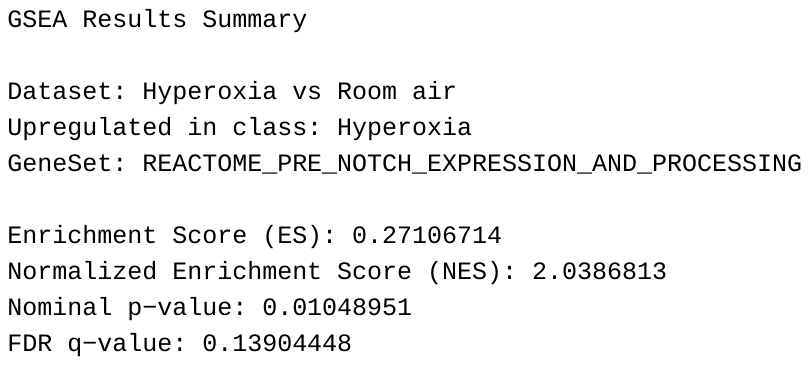

**FIGURE S6**

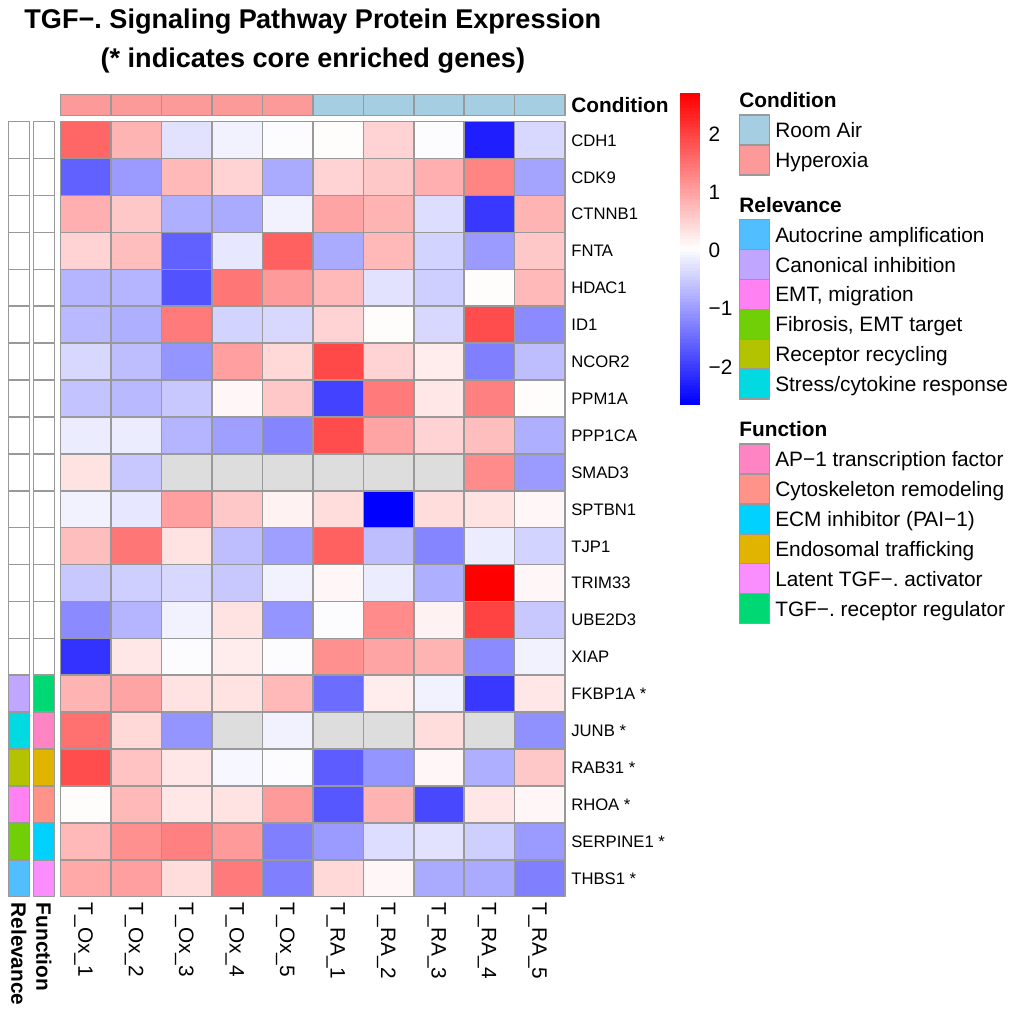

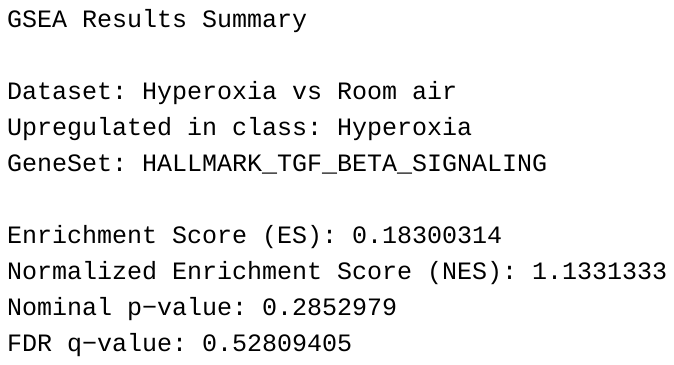
